# Ribosomal RNA Synthesis is a Lethal Vulnerability During Reductive Stress In *C*. *elegans*

**DOI:** 10.1101/2025.11.24.690266

**Authors:** Jen F. Rotti, Fasih Ahsan, Nicole L. Stuhr, Sinclair Emans, Alexander Soukas, Armen Yerevanian

## Abstract

Reductive stress has remained underappreciated as a significant disrupter of redox homeostasis. Recent studies have begun to link the accumulation of NADH and NADPH to the development and progression of metabolic diseases such as cancer, cardiac disease, and diabetes. Further research is needed to understand how cellular responses to reductive stress are governed. In this study we use the nematode *Caenorhabditis elegans* to examine the phenomenon of catastrophic reductive-death caused by combined biguanide treatment and *fasn-1* deficiency. This process of synergistic reductive stress correlates with aberrant alternations in nucleolar morphology. The absence of *fasn-1* activity blocks phenformin-mediated reduction in nucleolar size in the hypodermis, potentially resulting in enhanced translation. We find that loss-of-function and RNAi-based knockdown of the catalytic RNA exosome subunit *crn-3* significantly increases resistance to toxic reductive stress. Multiple other genes involved in rRNA synthesis recapitulate this phenotype. We postulate that this reversal of reductive death can be attributed to impaired ribosomal RNA biogenesis that promotes tolerance of the accumulation of reducing equivalents NADPH and NADH and preventing the accumulation of GSH. Overall, we identify a novel mechanism by which pathologic states of reductive stress-related diseases can be ameliorated.

## Introduction

Redox reactions are essential for the maintenance of organismal health via cell signaling, gene regulation, and metabolite generation (Xiao et al., 2018; Sies, Mailloux & Jakob, 2024). NADH/NAD+, NADPH/NADP+, and GSH/GSSG, are the three major electron carriers that act as substrates for mitochondrial and cytoplasmic redox reactions, powering critical biological processes (Xiao et al., 2018). An imbalance of oxidant and antioxidant species can lead to cellular dysfunction and adverse health outcomes (Xiao et al., 2018; Sies, Mailloux & Jakob, 2024). Redox signaling during development is particularly important as early life ROS signaling can cause epigenetic changes that promote stress resistance and longevity later in life (Bazopoulou et al., 2019). It is therefore important to understand the mechanisms through which redox stress can be mitigated.

Reductive stress – the imbalance of redox species in favor of an antioxidant state - has been shown to be involved in the pathology of several diseases. For example, *in vivo* administration of the antioxidants *N*-acetylcysteine or vitamin E accelerated the progression of lung cancer in mice possibly by preventing the activation of the tumor suppressor p53 which is typically activated by oxidative stress (Sayin et al., 2014). Overexpression of molecular chaperone CryAB causes redox stress that can accelerate cardiac hypertrophy but is reversed upon inhibition of glucose-6-phosphate dehydrogenase (G6pd). This links NADPH production via the pentose phosphate pathway to cardiac disease (Rajasekaran et al., 2011). In addition, increased glutathione peroxidase activity has also been shown to cause insulin resistance in mice (McClung et al., 2004). Thus, identifying ways to prevent reductive stress could provide crucial insight into disease pathology and potential therapeutic interventions.

Metformin, the most widely prescribed anti-type 2 diabetes medication, is a member of a class of drugs called biguanides that have also been shown to alter cellular metabolism, resulting in extended health span and lifespan in a wide variety of model organisms including invertebrate worms, mice, and non-human primates (Soukas, Hao & Wu, 2019; Yang et al., 2024). Investigation into the genetic mechanisms through which biguanides extend lifespan is ongoing.

Recent work in our lab highlights the importance of maintaining redox homeostasis upon biguanide administration. Our recent studies have suggested that phenformin, a potent sister biguanide to metformin, significantly alters redox homeostasis in the invertebrate nematode *C. elegans* by enhancing the reductive ratio of NADH/NAD+, NADPH/NADP+, and GSH/GSSG couples (Ahsan et al., 2025). We observed that inhibition of *de novo* lipogenesis, the major organismal fatty acid biosynthetic process, function as a critical brake to prevent unchecked reductive stress induced by multiple NADPH accumulating paradigms (Ahsan et al., 2025).

Fatty acid synthesis is the major consumer of NADPH within *C. elegans* and other eukaryotes. In the FASN-1-catalyzed conversion of malonyl-CoA into palmitate, 14 molecules of NADPH are oxidized into NADP+. Thus, the inhibition of essential *de novo* fatty acid synthesis genes *fasn-1* and *pod-2* results in a toxic accumulation of antioxidant species that is not rescued upon palmitate end-product supplementation (Ahsan et al., 2025). Supplementation with malic acid or glucose similarly promotes NADPH accumulation and phenocopies this synthetic lethal phenotype when combined with deficient fatty acid biosynthesis (Ahsan et al., 2025). To prevent reductive stress, *C. elegans* and mammalian cellular models tend to increase *de novo* fatty acid synthesis as a detoxifying strategy (Ahsan et al., 2025). These findings emphasize the importance of fatty acid synthesis in maintaining redox homeostasis when challenged with excessive reducing power caused by biguanides and other pharmacological and nutritional interventions. The pathology of phenformin-induced reductive stress remains unclear. In this paper, we aim to understand how organisms may protect themselves against reductive stress.

The pathology of phenformin-induced reductive stress remains unclear. In this paper, we aim to understand how organisms may protect themselves against reductive stress. Emerging studies have begun to recognize the role of the nucleolus in cellular responses to stress. This sub-nuclear compartment is essential to ribosome biogenesis and stores hundreds to tens of thousands of copies of ribosomal DNA (rDNA) depending on the species and is transcribed to generate ribosomal RNA (rRNA) that forms the structural core of ribosomes (González-Arzola, 2024). In response to starvation, rRNA synthesis is reduced in *C. elegans* and mammalian cell models as a way to potentially conserve energy (Shovon et al., 2025; Lu et al., 2025; Wu et al., 2018). Additionally, inhibiting RNA polymerase I activity has been shown to protect against toxicities associated with late-in-life administration of metformin in *C. elegans*, potentially by protecting metabolic plasticity and energetic homeostasis (Sharifi et al., 2024). Thus, targeting rRNA synthesis may aid in adaption to stress.

Performing a high-throughput RNAi genetic screen in *C. elegans*, we demonstrate that phenformin-induced reductive stress is associated with altered nucleolar morphology and that activation of a nucleolar stress response pathway increases tolerance of reducing equivalent accumulation. These findings highlight rRNA biogenesis as a lethal vulnerability during biguanide-induced reductive stress.

## Results

### RNAi Knockdown of *crn-3* and *bud-23* Protects Against Toxic Reductive Stress

We harnessed reverse genetics by combining RNA interference (RNAi) and auxin inducible degron (AID) technology with pharmacological interventions to examine factors that mitigate the toxicity of a highly reductive cellular environment. To spatiotemporally visualize and knockdown *fasn-1*, we crossed animals bearing a somatically expressed *Arabidopsis thaliana* auxin receptor transport inhibitor response 1 protein transgene [AtTIR-1(F79G)] with animals expressing an endogenously tagged N-terminal 3xFLAG::AID*::GFP transgene (Ahsan et al., 2025; Zhang et al., 2015).

AID activation via 5-phenyl-1H-indole-3-acetic acid (5-Ph-IAA) administration was used to depleted FASN-1 protein levels. Correspondingly, survival was significantly reduced with approximately 20% of reached adulthood after 120 hours when treated with the drug from L1 of hatching (Figure 1 C). When 5-Ph-IAA treatment was combined with a lifespan-extending dose of 4.5 mM phenformin from L1 of hatching, a near fully penetrant larval lethal phenotype was observed with less than 5% of animals reaching adulthood (Figures 1C). These results were consistent with the reductive death phenotype observed upon combining *fasn-1* RNAi with phenformin and other NADPH promoting interventions such as malic acid and glucose supplementation (Ahsan et al., 2025).

**Figure 1:**
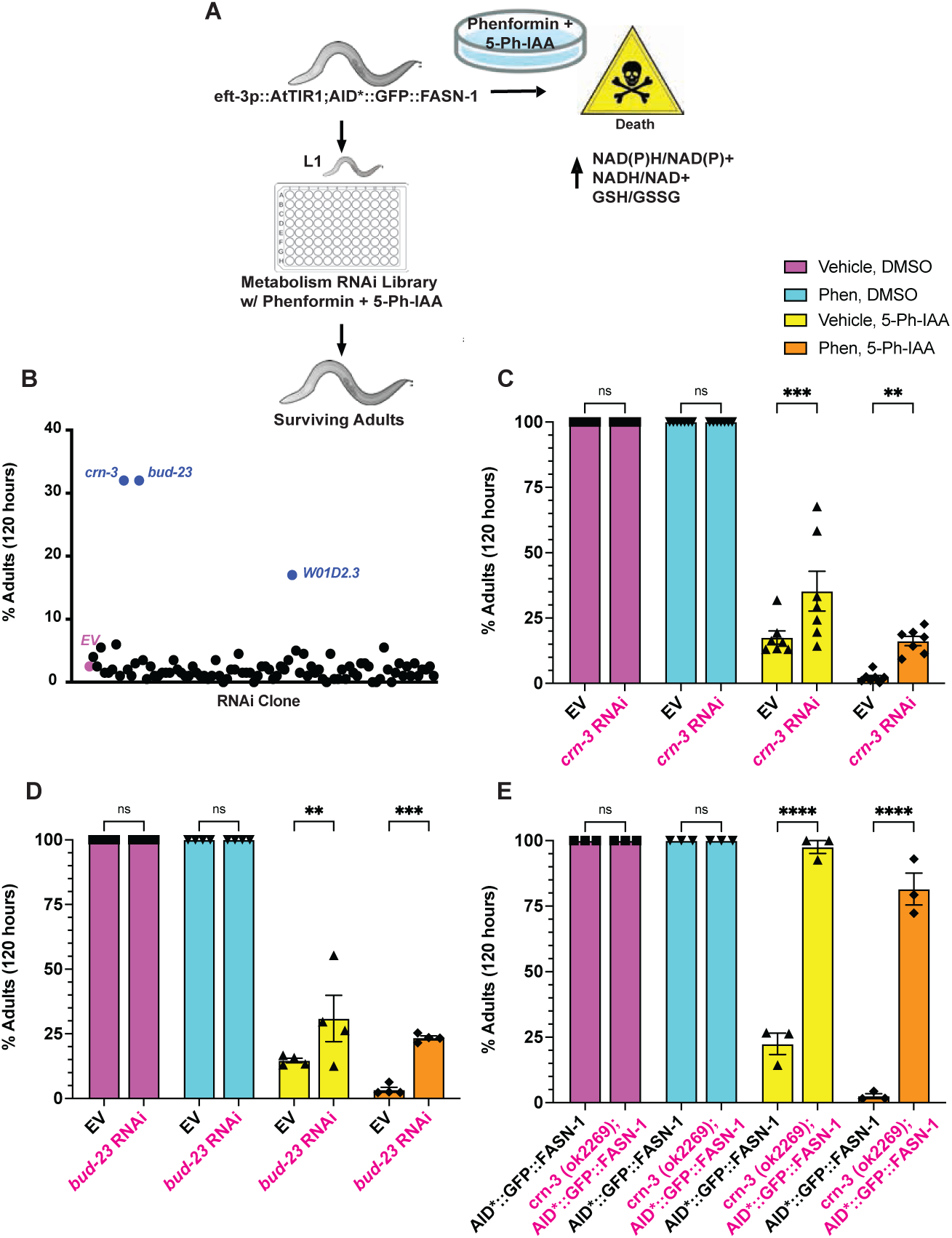
Reverse Genetic Screen Identifies *crn-3* and *bud-23* RNAi as Suppressors of Biguanide-Induced Toxicity in the Absence of *de Novo Fatty Acid Synthesis*. (A) Reverse genetic metabolic RNAi screen assay experimental design (B) Secondary retesting of 91 RNAi clones of interest from initial screen. (C-E) Quantification of developmental assays on (C) EV vs *crn-3* RNAi (D) EV vs *bud-23* RNAi and (E) control animals vs *crn-3(ok2269* mutants. 3≥_biological replicates in technical triplicate were performed. n = 200 animals per plate. All error bars represent mean +/- SEM unless otherwise noted. Significance was assessed using a two-way ANOVA with Sidak correction of multiple comparison testing. ns p > 0.05, * p < 0.05, ** p < 0.01, *** p < 0.001, **** p < 0.0001.

Leveraging this AID system in tandem with RNA interference (RNAi) technology, we sought to uncover epistatic genetic interactions that underlie this synergistic death response. We performed a high throughput genetic screen, treating L1 stage *fasn-1* deficient nematodes with phenformin and concomitant feeding of an established RNAi library targeting 1046 metabolism associated genes (Figure 1A)(Qin et al., 2025). From this initial screen, we identified two genes – *crn-*3/CeExosc-10 and *c27f2.4/*CeBUD*-23* - whose loss significantly improved survival (Figure 1B).

We validate RNAi knockdown efficiency for *crn-3* and *bud-23* via qRT-PCR (Figure S1 F). RNAi of these genes resulted in improved organismal survival in the face of *fasn-1* deficiency (Figure 1 C-D; Figure S1 C). To corroborate our findings genetically, we crossed AID*::GFP:: FASN-1 animals with a *crn-3(ok2269)* null mutant which has a 1049 bp deletion within the coding sequence. We observed that *crn-3* mutant worms exhibited delayed development - taking approximately 24 hours longer to reach the L4 stage compared to wild-type animals - consistent with hypomorphic phenotypes seen with impaired ribosome biogenesis (Figure S1 D-E). Notably, despite this significant delay in development, *crn-3* mutants exhibited a complete restoration of survival to adulthood compared to wild-type animals after 72 to 120 hours of 5-Ph-IAA treatment alone (Figure 1E; Figure S1D-E). Strikingly, the majority of *crn-3* mutants treated with both 5-Ph-IAA and phenformin reached adulthood after 120 hours (Figure 1E).

To rule out loss of AID efficiency as a cause for the observed increased stress resistance of *crn-3* mutants, we evaluated FASN-1 expression upon combined inactivation and *crn-3* null mutation. Immunoblotting confirmed that genetic ablation of crn-3 did not alter FASN-1 depletion efficacy, and moreover revealed that *crn-3* mutant animals overexpress FASN-1 protein compared to the control, implicating *crn-3* as a possible negative regulator of *fasn-1* (Figure S1 A-B).

### Mutation of crn-3 Protects Against Accelerated Death Associated with Post-Developmental Phenformin and *fasn-1* RNAi Treatment

To determine if *crn-3* and *bud-23* deficiency was sufficient to protect against reductive stress throughout organism lifespan, we performed longitudinal survival analyses. We observed that that *crn-3* null mutants were naturally short lived relative to wild-type animals (Figure 2 A-C; Table S1). Consistent with our expectations, *crn-3* mutants experienced a significantly delayed onset of death when compared to wild type worms treated with *fasn-1* RNAi and phenformin post-developmentally from L4 of hatching (figure 2 A-C; Table S1). In contrast, *bud-23* mutant animals did not delay the onset of this synthetic death phenotype (Figure S2A-C; Table S2). Both the *crn-3* and *bud-23* mutants exhibited impaired phenformin-mediated lifespan extension compared to wild-type (Figure 2A-C; Figure S2A-C; Table S1; Table S2, suggesting that that function of these two factors may be required for phenformin-lifespan extension, but are dispensable to prevent accelerated death in the context of *fasn-1* deficiency.

**Figure 2.**
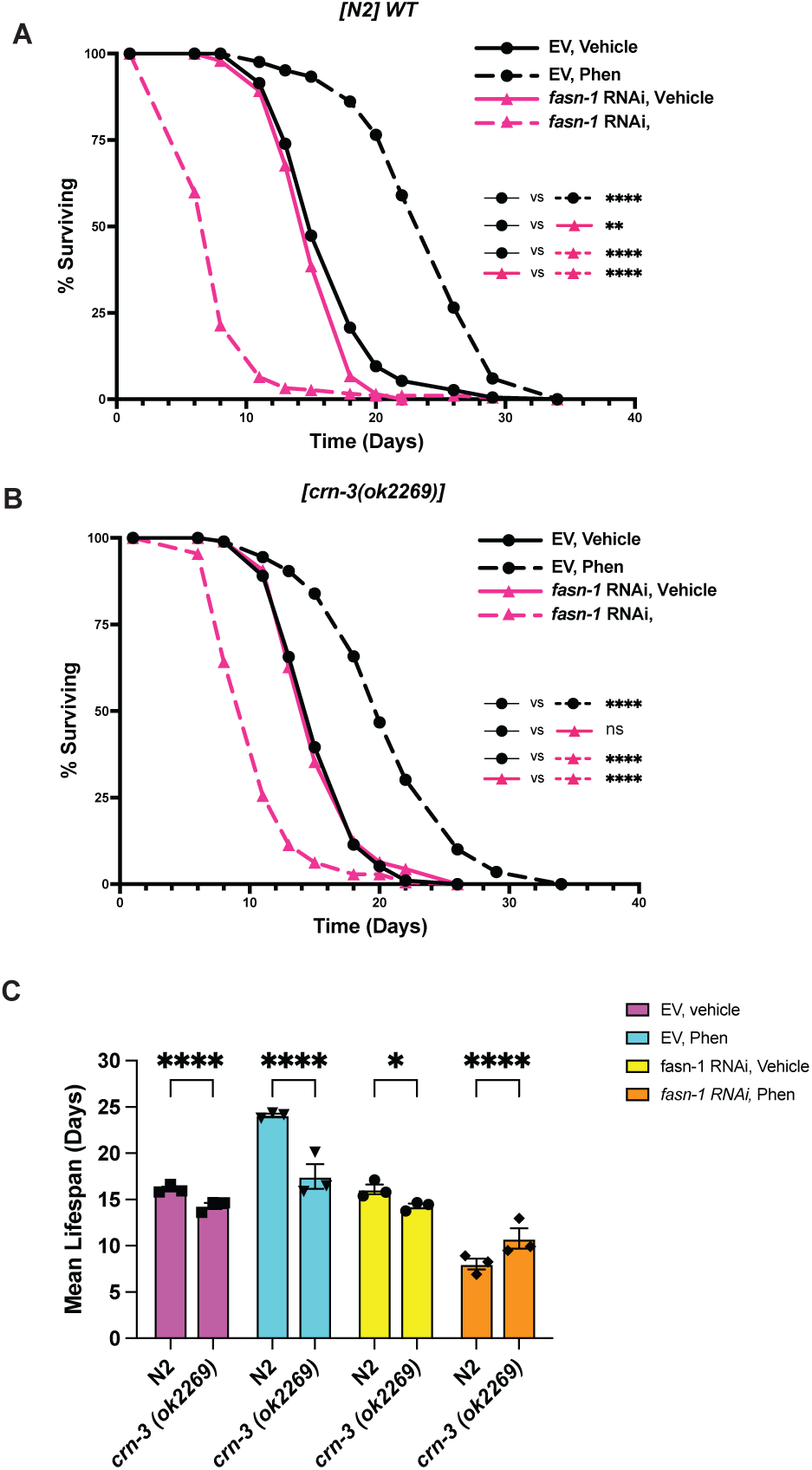
Genetic Mutation of *crn-3* Delays Onset of Accelerated Death In *fasn-1* Deficient Animals Treated with Phenformin. (A-B) Lifespan analyses of (A) wild-type and (B) *crn-3(ok2269)* animals fed EV or *fasn-1* RNAi and treated with either vehicle or 4.5 mM phenformin from L4 of development (C) Visulaization of mean lifespan results for each condition in Figure 2A-B. Three independent biological replicates were performed in triplicate n = 3. All mutant animals were backcrossed 4-6 times to wild-type animals to remove potential background mutations. Significance was assessed in (A-B) using log rank analyses. ns p > 0.05, * p < 0.05, ** p < 0.01, *** p < 0.001, **** p < 0.0001. See Table S1 for log rank analyses.

### FASN-1 is Necessary for Phenformin-Mediated Regulation of Nucleolar Size and Translational Control

The observation that the knockdown of *crn-3* and *bud-23* – two genes involved in rRNA biogenesis - significantly improved survival prompted an investigation into the effects of phenformin-induced reductive stress on rRNA synthesis. We hypothesized that an overexpression of rRNA could be a pathological consequence of reductive death. To test this, we performed experiments to assess nuclear morphology using a translational reporter for the nucleolar protein fibrillarin (ie. *fib-1)* with co-staining of nuclear DNA (Figure 3A). We used nucleolar size as an indicator of nucleolar activity given that enhanced rDNA transcription and carcinogenesis are positively correlated with larger nucleoli, whilst small nucleoli are reciprocally predictive of pro-longevity response (Tiku et al., 2017; Ma et al., 2016). Animals were assessed on day 2 of adulthood just as the first couple of worms were starting to die on plates treated with phenformin and *fasn-1* RNAi, signaling the onset of death. We observed that a lifespan extending dose of 4.5 mM phenformin significantly reduced the nucleolar to nuclear area ratio in posterior hypodermal cells imaged in the tail region. Notably, this effect was totally reversed upon RNAi knockdown of *fasn-1*, suggesting that *fasn-1* may act as a negative regulator of rRNA synthesis (Figure 3B).

**Figure 3.**
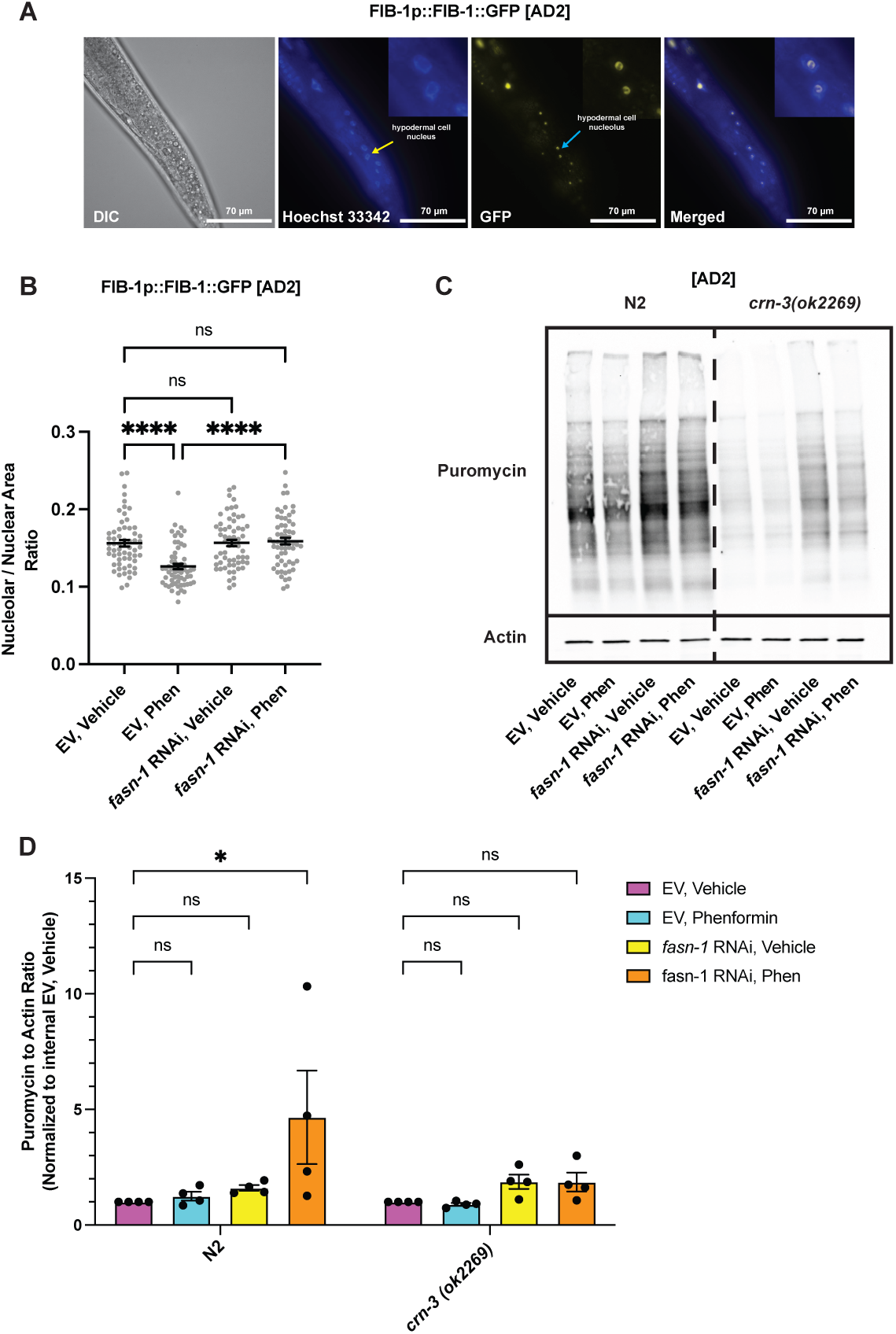
*f*asn-1 Deficiency Blocks Phenformin-Mediated Regulation of Nucleolar Size and Translation. (A) Representative images of posterior hypodermal cells. Cells were imaged at 63x magnification under a GFP filter to detect *fib-1p::fib-1::GFP*. Worms were additionally stained with Hoechst 33342 to visualize nuclear DNA and imaged under a BFP filter. (B) Quantification of nucleolar to nuclear ratio in animals treated with vehicle or 4.5 mM phenformin with or without feeding of *fasn-1* RNAi from L4 to Day 2 of adulthood (AD2). Data is representative of three independent biological replicates with n = 20 worms per condition. (C) Representative SUrface Sensing of Translation (SUnSET) puromycin incorporation immunoblotting in wildtype or *crn-3(ok2269)* animals treated with vehicle or 4.5 mM phenformin with or without *fasn-1* RNAi from L4 to AD2. (D) Quantification of puromycin incorporation immunoblots. Puromycin incorporation assays were performed in 4 independent biological replicates n = 4. Due to the developmental delay in *crn-3(ok2269)* mutants, puromycin incorporation assays were performed a day after the wildtype control. All error bars represent mean +/- SEM unless otherwise noted. Significance was assessed in using a one-way ANOVA in (B) and a two-way ANOVA in (E) with Sidak correction of multiple comparison testing. ns p > 0.05, * p < 0.05, ** p < 0.01, *** p < 0.001, **** p < 0.0001.

Inhibition of hypodermal nucleolar activity and ribosome biogenesis has been implicated in pro-longevity responses and stress resistance (Tiku, et al., 2017; Dalton & Curran, 2017; Zhao et al., 2023). RNA sequencing revealed that transcripts upregulated upon combined phenformin, and *fasn-1* RNAi treatment were predicted to be expressed in the hypodermis or epithelial system (Figure S3 A). Additionally, lifespan analysis revealed that hypodermal but not intestine specific RNAi knockdown of *fasn-1* phenocopied the accelerated demise observed upon tandem whole-organism RNAi and phenformin treatment (Figure S3 B-D). Together this suggests that the hypodermis may play an especially important role in the defense against reductive stress. Thus, we focused on the nucleolar morphology of hypodermal cells.

Animals were assessed on day 2 of adulthood which is when the first worms start to die on the combined phenformin and *fasn-1* RNAi condition. We observed that a lifespan extending dose of 4.5 mM phenformin significantly reduced the nucleolar to nuclear area ratio in hypodermal cells imaged in the tail region. Notably, this effect was totally reversed upon RNAi knockdown of *fasn-1*, suggesting that *fasn-1* may act as a negative regulator of rRNA synthesis (Figure 3B).

We wondered whether enhanced nucleolar activity might cause an increase in the downstream translation of proteins. Puromycin incorporation assays revealed that *fasn-1* RNAi may enhance nascent protein synthesis in both wildtype and *crn-3* mutant animals, although it was not statistically significant (Figure 3C-D). Notably, the combination of *fasn-1* RNAi and phenformin significantly increased translation in wildtype animals but not *crn-3* mutants (Figure 3D). This could implicate aberrant translation in the pathology of phenformin-induced reductive death, that may be reversed by inhibiting the activity of *crn-3*.

### Enhanced rRNA biogenesis Phenocopies Phenformin - Induced Hypodermal Stress Response in the Absence of FASN-1 Activity

As previously suggested, increased expression of hypodermal genes occurs upon tandem phenformin and *fasn-1* RNAi treatment (Figure S3 A). Knockdown of *fasn-1* has previously been shown to trigger the induction of hypodermal stress response gene *nlp-29* (Lee et al., 2010). We thus hypothesized that biguanide treatment synergistically enhances the primed epidermal stress state in *fasn-1* deficient animals. RNA sequencing revealed that phenformin and *fasn-1* RNAi synergize to drive elevated *nlp-29* expression (Figure 4A). We verified these results at the protein level, using an integrated *nlp-29p:GFP;col-12p::dsRed* translational reporter treated with combined *fasn-1* RNAi and phenformin (Figure 4B) or when crossed with *fasn-1(fr8)* hypomorphic animals (Figure 4 E-F). Notably the temporal expression of *nlp-29* was closely associated with the mean timepoint of death for each condition (Figure 2A; Figure 4B). Combined, these data suggest that aggravated hypodermal induction of *nlp-29* expression acts as a biomarker of biguanide-induced reductive death upon *fasn-1* deficiency.

**Figure 4.**
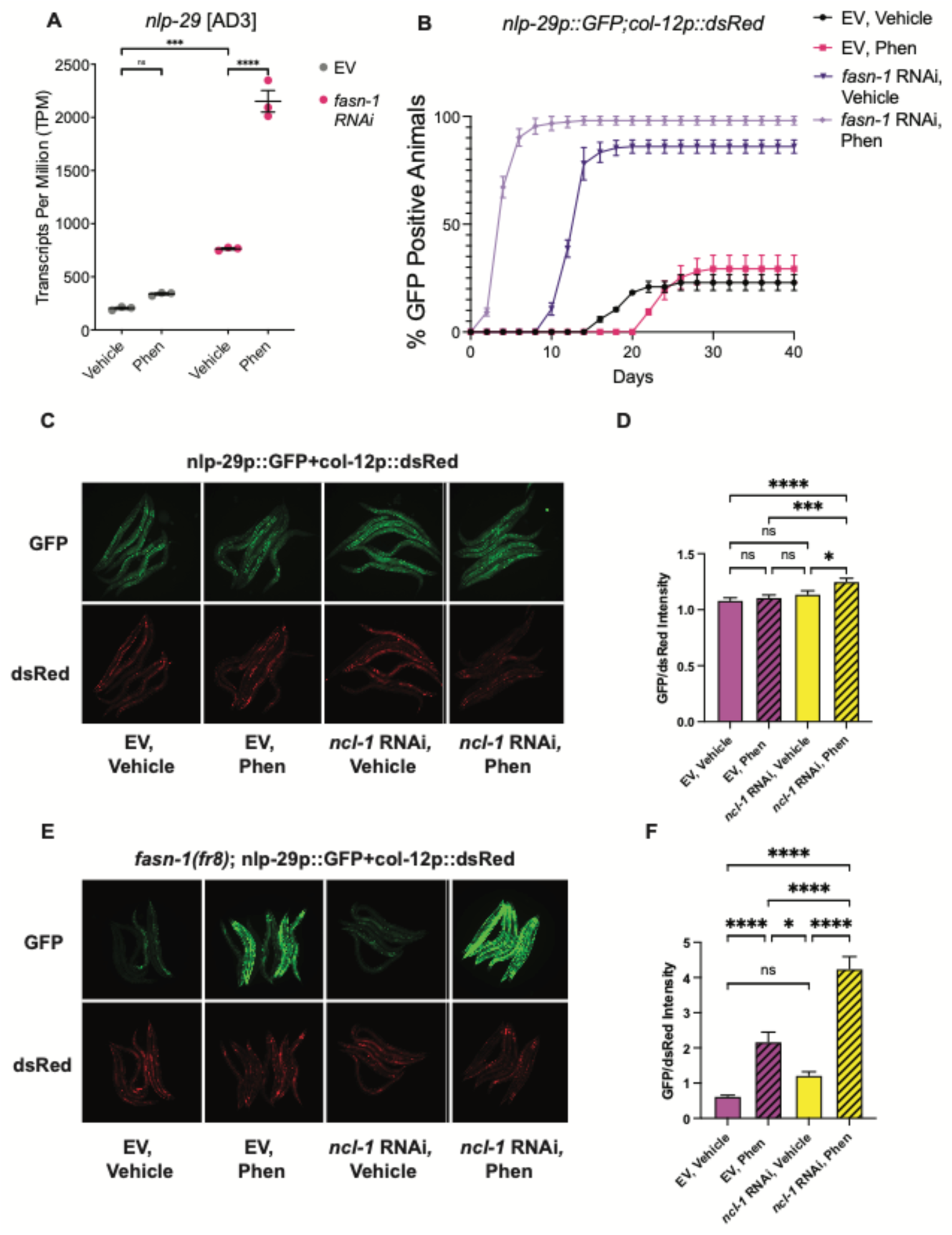
*ncl-1* RNAi Synergistically Activates Hypodermal Stress Gene *nlp-29* in Combination with Phenformin of FASN-1 Depletion. (A) Abundance of *nlp-29* mRNA in N2 wildtype worms fed with EV or *fasn-1* RNAi and treated with vehicle or 4.5 mM phenformin. (B) Quantification of GFP-positive *nlp-29p::GFP;col-12p::dsRed* animals over time treated with or without *fasn-1* and 4.5 mM phenformin. (C-D) Representative images and quantification of fluorescence in *nlp-29p::GFP;col12p::dsRed* animals with or without *ncl-1* RNAi and 4.5 mM phenformin from L4 to AD2. (E-F) Representative images and quantification of fluorescence of *fasn-1(fr8); nlp-29p::GFP;col12p::dsRed* animals treated with or without *ncl-1* RNAi and 4.5 mM phenformin from L4 to AD2. (A,B,D,F). A total of 150 worms were imaged per condition, across three biological replicates. All error bars represent mean +/- SEM unless otherwise noted. Significance was assessed in (D,F) using a one-way ANOVA with Sidak correction of multiple comparison testing. ns p > 0.05, * p < 0.05, ** p < 0.01, *** p < 0.001, **** p < 0.0001.

Given our earlier results suggesting that *fasn-1* may be a negative regulator of nucleolar size, we wondered whether the enhancing nucleolar activity could phenocopy the induction or *nlp-29* caused by reductive stress when combined with phenformin or *fasn-1* deficiency. We thus leveraged RNAi inactivation of *ncl-1,* a histone chaperone protein that is a known suppressor of nucleolar size and rRNA biogenesis associated with accelerated aging (Tiku et al., 2017). Consistent with our hypothesis, we observed a slight, yet significant, activation of the hypodermal stress response when *ncl-1* RNAi was combined with phenformin treatment (Figure 4D). Compellingly, we observed that combining deficiency in *fasn-1* and *ncl-1* with phenformin resulted in synergistic activation of *nlp-29* to a degree that exceeded all other experimental groups (Figure 4 E-F). These findings suggest that enhancing rRNA biogenesis may increase sensitivity to reductive stress caused by biguanides and *fasn-1* deficiency.

### Disruption of rRNA Transcription, Modification, and Processing Protects Against Reductive Stress

Given that the canonical functions of *crn-3* and *bud-23* converge upon rRNA biogenesis and that enhancing rRNA transcription sensitized animals the effects of phenformin and *fasn-1* deficiency, we speculated that perturbing rRNA biosynthesis would prevent reductive death. The production of functional rRNA is a highly complex process that involves many different of post-transcriptional modifications and translocation from the nucleolus to the cytoplasm. We were interested to see whether impairing different steps in RNA polymerase I-mediated rRNA synthesis improved survival under reductive stress conditions during development. We first inhibited rRNA synthesis at its source by targeting RNA polymerase I transcription initiation factor *tif-1a/*CeRRN-3. The results from our developmental assays revealed that *tif-1a* RNAi significantly increased the number of surviving *fasn-1* deficient animals with and without phenformin. Similarly, RNAi knockdown of the nuclear rRNA processing genes *t22h9.1/*CeRRP*-36* and *lpd-6/* PPAN-P2RY11/CePPAN-P2RY11, the exportin *xpo-1* which facilitates the exportation of rRNA transcripts and ribosomal subunits from the nucleus into the cytosol for downstream ribosome assembly, and *riok-1*/CeRIO1 which facilitates the maturation of the 18S rRNA in the cytoplasm all significantly reversed reductive death. (Dörner et al., 2023)(Figure 5A-F; Figure S4). These results suggest that disrupting rRNA biogenesis at various points in the pathway, from early-stage transcription to nuclear processing, to nuclear exportation, to late-stage cytoplasmic maturation can activate a nucleolar stress response that, in turn, promotes resistance to reductive stress.

**Figure 5.**
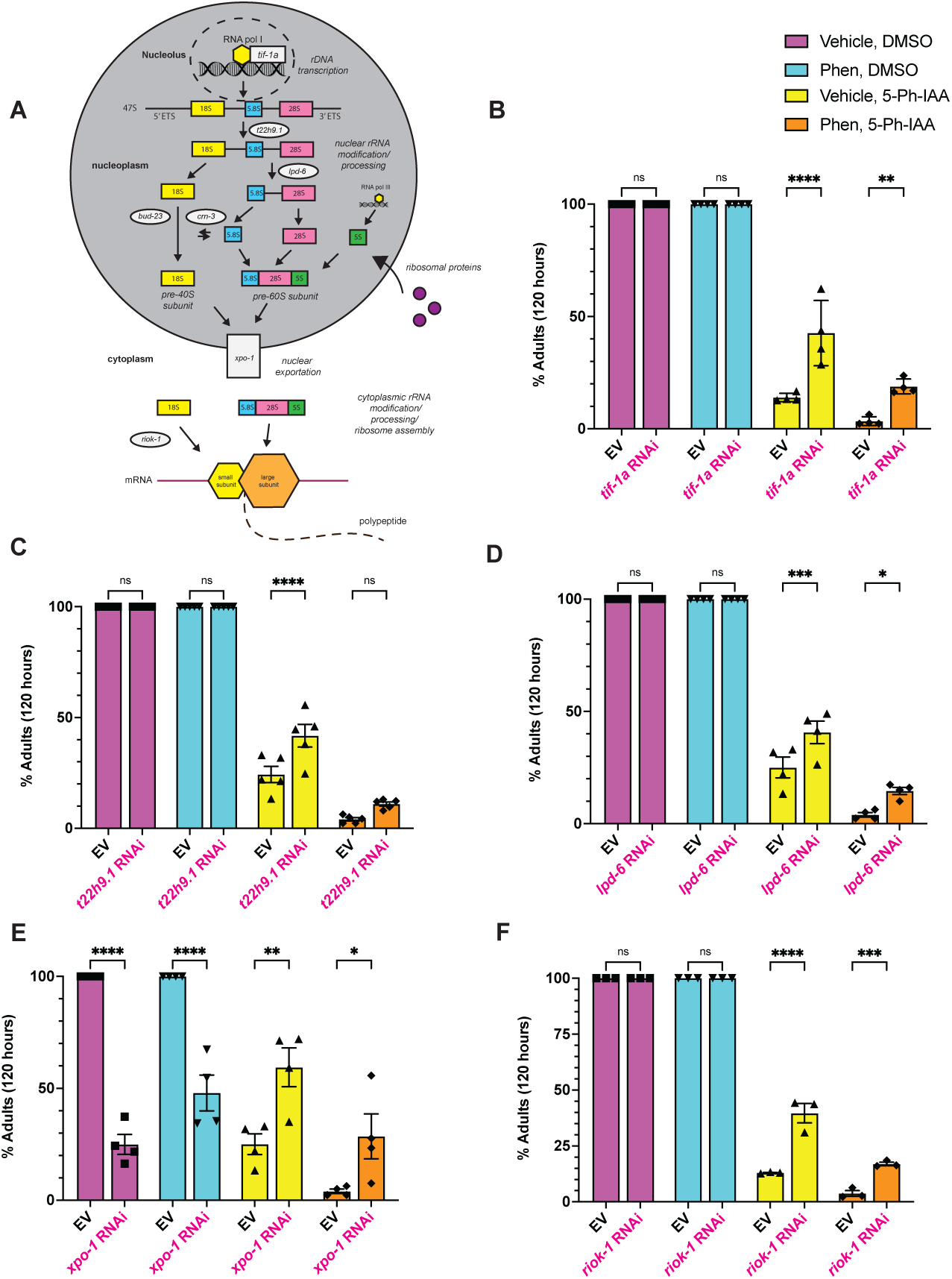
Inhibition of rRNA Biogenesis Significantly Reverses Toxicity. (A) Simplified schematic of RNA polymerase I-mediated rRNA synthesis pathway. RNA polymerase I binds to ribosomal DNA in the nucleolus and transcribes the tri-cistronic 47S pre-rRNA transcript. Subsequent cleavage, modification, and exportation of the rRNA from the nucleus into the cytoplasm for final processing occurs to produce mature 18S rRNA, 5.8S rRNA, and 28S rRNA. Together with 5S rRNA which is transcribed separately by RNA polymaser III, the small and large subunits are assembled. (B-F) Quantification of development assay in animals were treated with various RNAi targeting genes involved in rRNA biogenesis. Experiments were performed in > 3 biological replicates in technical triplicate. n = 200 animals per plate. All error bars represent mean +/- SEM unless otherwise noted. Significance was assessed in (D-E, H) using a two-way ANOVA with Sidak correction of multiple comparison testing. ns p > 0.05, * p < 0.05, ** p < 0.01, *** p < 0.001, **** p < 0.0001.

### Inhibiting *crn-3* Increases Tolerance of NADH Reductive Stress and Increases Oxidized Glutathione Pools

Previous work done by our lab suggests that the accumulation of antioxidant species is the likely cause of accelerated death upon combined phenformin and *fasn-1* RNAi (Ahsan, 2025). This led us to investigate whether impairment of RNA exosome or rRNA biogenesis could directly mitigate reductive stress and improve survival outcomes. We conducted *ex vivo* ratiometric fluorescence assays to investigate redox dynamics in wild-type versus *crn-3*(ok2269) mutants.

We found that overall NADPH/NADP+ and NADH/NAD+ ratios were elevated in *crn-3* mutants when compared to wildtype animals across all treatment groups (Figure 6 A-B). We were interested in whether *crn-3* mutants responded differently than wild-type animals when challenged by reductive stressors. Thus, we looked at the general trends in redox ratios within each strain. In both *crn-3* and wildtype animals, the general trend of NADPH/NADP+ ratios increased with phenformin, *fasn-1* RNAi, and the combination of both, consistent with the accumulation of NADPH described in Ahsan et al. (2025).

**Figure 6.**
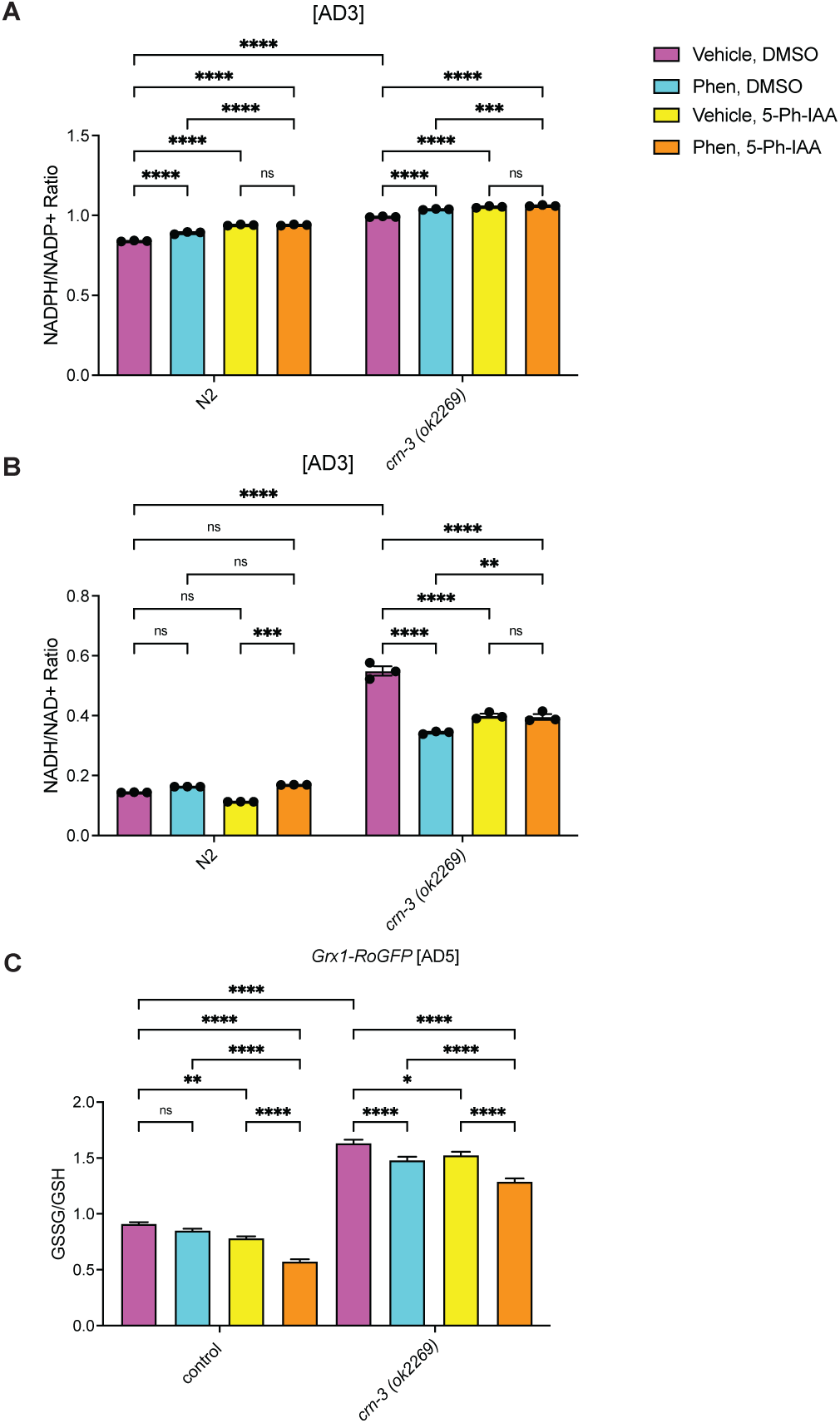
Inhibition of *crn-3* via Genetic Mutation Alters NADPH/NADP+, NADH/NAD+, amd GSSG/GSH Ratios. (A-B) Quantification of the NADPH/NADP+ (A) and NADH/NAD+ (B) ratios in N2 wild-type and *crn-3(ok2269)* animals treated with or without *fasn-1* RNAi and 4.5 mM phenformin from L4 of development to AD3. Approximately 1000 animals were harvested for each condition per replicate. Due to the developmental delay in *crn-3(ok2269)*, redox measurements were performed a day after wild-type in order for stage-matching of strains. Three independent biological replicates were performed n=3 (C) Fluoresecent ratiometric expression of *rpl-17p::Grx1-roGFP2* in control and *crn-3(ok2269)* animals treated with *fasn-1* RNAi or 4.5 mM phenformin from L4 to AD5. A total of 30 animals were imaged per condition. Experiment was performed in two biological replicates n=2. All error bars represent mean +/- SEM unless otherwise noted. Significance was assessed in (A-D) using a two-way ANOVA with Sidak correction of multiple comparison testing. ns p > 0.05, * p < 0.05, ** p < 0.01, *** p < 0.001, **** p < 0.0001.

We observed that phenformin alone caused a non-significant increase in the NADH/NAD+ ratio while *fasn-1* RNAi caused a non-significant decrease in the NADH/NAD+ ratio in wild-type animals (Figure 6B). The combination of both treatments resulted in a significant increase in the NADH/NAD+ ratio compared to *fasn-1* RNAi alone, restoring the NADH/NAD+ ratio to baseline (Figure 6B). This suggests that phenformin and *fasn-1* synergize, causing the accumulation of NADH in the *fasn-1* deficient animals.

Meanwhile, results measuring the NADH/NAD+ ratio in *crn-3* mutants revealed a very different pattern. Phenformin caused a significant decrease in the NADH/NAD+ ratio, as did *fasn-1* RNAi alone (Figure 6B). Notably, combination of the two treatments did not result in a synergistic elevation of the NADH/NAD+ ratio (Figure 6B). Together, these results suggest that *crn-3* mutation may increase tolerance to high levels of reducing equivalents caused by phenformin and *fasn-1* deficiency rather than directly preventing their accumulation. However, it is also notable that the of effects of phenformin and *fasn-1* RNAi on NADH/NAD+ ratio dynamics vary significantly between strains, suggesting that *crn-3* deficiency can fundamentally alter cellular response to exogenous reductive stress caused by phenformin.

To determine whether inhibition of *crn-3* prevented GSH reductive stress, we crossed *crn-3(ok2269)* animals with an *in vivo* GSH/GSSH reporter. We observed a synthetic decrease in GSSG/GSH ratios upon combined phenformin and *fasn-1* RNAi in the control strain, consistent with GSH reductive stress (Figure 6C). Notably, GSSG/GSH ratios were significantly increased in *crn-3* mutant worms across all treatment conditions when compared to control animals (Figure 6C). However, a similar trend of a synergistic decrease in GSSG/GSH ratios was still observed in *crn-3* animals treated with combined phenformin and *fasn-1* RNAi (Figure 6C). Together is suggests that *crn-3* mutation shifts the glutathione redox pools towards a more oxidized state, potentially alleviating toxic GSH reductive stress.

### Protective Effects of Inhibiting rRNA Biogenesis Against Reductive Death are *rsks-1* Dependent

Given our observations that combined *fasn-1* RNAi and phenformin treatment resulted in a possible increase in global translation, we hypothesized that aberrant increases in translation could also be a potential cause of the accelerated reductive-death phenotype. We reasoned that if inhibition of rRNA biogenesis protects against reductive death, then inhibiting translation by targeting downstream ribosomal activity would have the same effect.

To answer this question, we treated *eft-3p::AtTIR1(F79G)::mRuby; 3xFLAG::AID*::GFP::FASN-1* animals with RNAi targeting *rsks-1*– the sole *C. elegans* homolog for human ribosomal protein S6-Kinase – thus impairing ribosomal translation initiation (Pan et al., 2007). As expect, we found that partial knockdown of *rsks-1* via RNAi significantly rescued the toxicity caused by 5-Ph-IAA treatment with or without phenformin (Figure 7A). To determine whether inhibition of *crn-3* or *tif-1a* protects against reductive death by impairing downstream ribosomal activity we crossed our *AID*::GFP::FASN-1* into a null mutant strain and treated them with *crn-3* of *tif-1a* RNAi. We hypothesized that the *rsks-1(ok1225)* allele and the *crn-3* or *tif-1*a RNAi would non-additively improve survival outcomes.

**Figure 7.**
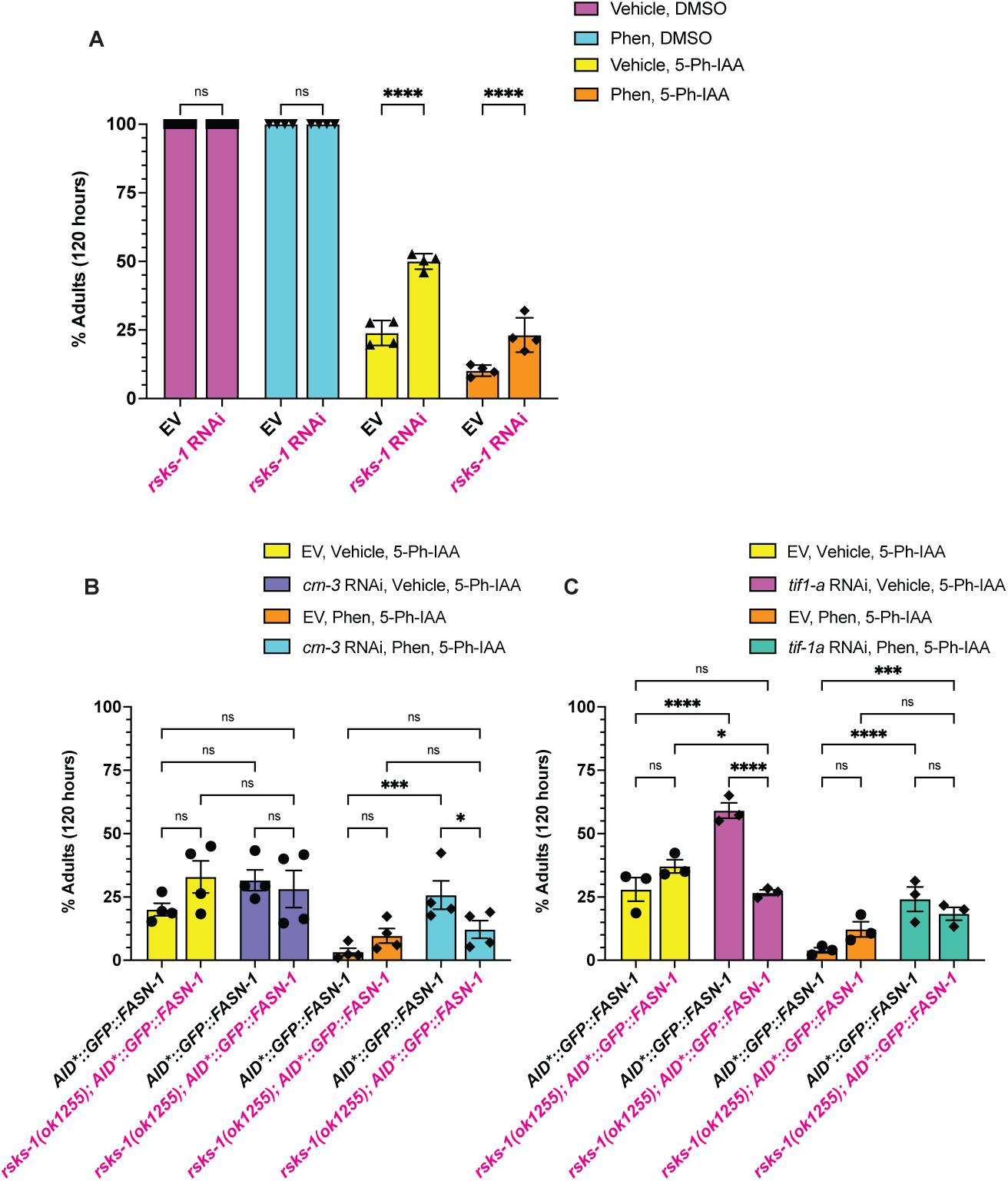
Ribosomal Kinase *rsks-1* Inhibition Protects Against Reductive Death, but is Necessary for rRNA - Mediated Reductive Stress Resistance. (A) Larval development assay performed on *eft-3p::AtTIR-1(F79G)::mRuby;3xFLAG::AID*::GFP::FASN-1* animals treated with either EV or *rsks-1* RNAi, vehicle or 4.5 mM phenformin, and either 0.1% DMSO or 5 µM 5-Ph-IAA. (B-C) Larval development assay performed on *eft-3p::AtTIR-1(F79G)::mRuby;3xFLAG::AID*::GFP::FASN-1* and *rsks-1(ok1255); eft-3p::AtTIR-1(F79G)::mRuby;3xFLAG::AID*::GFP::FASN-1* animals. Worms were fed either an EV control or RNAi and treated with a water vehicle or 4.5 mM phenformin with or without 0.1% DMSO or 5 µM 5-Ph-IAA. All animals grown on 0.1% DMSO controls reached adulthood after 120 hours, but are not shown. Results for (A-C) were compiled from 3-4 independent biological replicates in technical triplicate where n=200 animals per plate. All error bars represent mean +/- SEM unless otherwise noted. Significance was assessed in (D-E, H) using a two-way ANOVA with Sidak correction of multiple comparison testing.

We found that the *rsks-1* null mutation alone did not significantly improve survival outcomes of animals treated with 5-Ph-IAA with or without phenformin (Figure 7B-C). As expected, *crn-3* and *tif-1a* RNAi significantly rescued lethality in control animals treated with 5- Ph-IAA and phenformin (Figure 7B-C). Counter to our expectations, *rsks-1* mutants fed *crn-3* or *tif-1a* RNAi exhibited worse survival outcomes than control animals, reversing the benefits of impaired rRNA biogenesis (Figure 7B-C). This could suggest that while direct inhibition of translation efficiency via *rsks-1* knockdown is sufficient to prevent toxicity, *crn-3* and *tif-1a* inhibition may act through a parallel pathway that is partially dependent upon *rsks-1* activity.

## Discussion

In this study we discovered that deficiency in a number of rRNA transcription and maturation factors averts biguanide-induced reductive death in *C. elegans*. Meanwhile, enhancing rRNA synthesis synergizes with *fasn-1* deficiency to aggravate death. Notably, *crn-3* deficiency significantly altered redox ratios, rewiring the worm’s response to the accumulation of reducing equivalents.

The reducing equivalent NADH is closely tied to cellular energetics through mitochondrial activity via complex I of the electron transport chain (ETC). Complex I is a NADH:uniquinone oxidoreductase that uses NADH as a source of electrons needed for oxidative phosphorylation that produces adenosine triphosphate (ATP) (). In times of high energetic demand, increased flux through the ETC may decrease mitochondrial NADH pools (). Conversely, low energetic demand and reduced mitochondrial activity or mitochondrial dysfunction levels may cause NADH levels to increase ().

We observed elevated basal NADPH/NADP+ and NADH/NAD+ ratios in *crn-3* mutants compared to wildtype animals. This could be consistent with the role of *crn-*3 in ribosomal biogenesis. It is estimated that 80% of all RNA species produced in the cell are rRNAs (). Thus, inhibiting rRNA synthesis could reduce energetic demand thus causing NADH levels to rise. Notably, this increase in NADH levels did not lead to reductive death in *crn-3* mutants, even in animals treated with phenformin and *fasn-1* RNAi. This suggests that inhibiting *crn-3* and potentially rRNA synthesis improves survival outcomes not by preventing the accumulation of reducing equivalents, but rather by activating downstream signaling pathways that promote tolerance to NADH reductive stress. However, Further investigation is needed to identify which genes and signaling pathways are activated in order to prevent reductive death.

In our investigation, we observed that phenformin administration caused distinct changes in nucleolar morphology. Phenformin alone significantly reduced the ratio of nucleolar to nuclear size in hypodermal cells. However, this was completely reversed in the absence of *fasn-1,* causing an increase in downstream translation. The discovery that inhibition of *fasn-1* may enhance rRNA biogenesis when combined with phenformin was intriguing given the fact that biguanides may promote longevity by decreasing global translation (Ahsan et al., 2025). These findings point to decreased rRNA synthesis as a possible mechanism through which biguanides inhibit translation.

To our knowledge, this is the first paper to identify biguanides and *fasn-1* as potential regulators of nucleolar size and activity in *C. elegans*. In MCF-7 breast cancer cells, metformin reduces rRNA transcription by activating AMPK and inhibiting succinate production via the TCA cycle to epigenetically regulate rDNA expression (Tanaka et al., 2019). Biguanide-mediated activation of AMPK has also been shown to increase expression of nucleolin and prevent nuclear localization of fibrillarin, proteins that are known to be negative and positive regulators of rRNA synthesis respectively (Kodiha et al., 2014). It has also been suggested that AMPK can directly phosphorylate *tif-1a* to suppresses its activity (Hoppe et al., 2009). Interestingly, our previous work shows that loss of *fasn-1* activity prevents phenformin-mediated phosphorylation of *aak-1* – the *C. elegans* homolog of human AMPK (Ahsan et al., 2025). This seemingly correlates with the prevention of nucleolar size reduction in *fasn-1* deficient animals treated with biguanide. Thus, future work could investigate whether phenformin-mediated regulation of nucleolar size relies upon a conserved AMPK dependent pathway in *C. elegans*.

Other groups have linked cellular energetics to rRNA synthesis via eNOSC (energy-dependent Nucleolar Silencing Complex) which is partially comprised of the NAD+ dependent histone deacetylase SIRT1. Thus, in cases of low NAD+ levels, such as glucose deprivation, eNOSC epigenetically silences rDNA transcription (Murayama et al., 2008). This work establishes a potential link between NADH reductive stress and enhanced ribosomal biogenesis where low NAD+ levels prevent eNOSC-mediated epigenetic silencing of rDNA. Thus, multiple energy sensing pathways may play a key role in regulating rRNA synthesis during phenformin-induced reductive stress.

The mechanisms through which inhibition of *fasn-1* increases nucleolar activity and translation in *C. elegans* is also unknown. Studies in cellular models have identified the mammalian homolog *FASN* as a regulator of rRNA biogenesis and nucleolar size. It has been proposed that *fasn-1* inhibition causes the accumulation of the intermediate metabolite malonyl-CoA, promoting the malonylation of lysine residues on histones and nucleolin (Sun et al., 2024; Zhang et al., 2023). As a result of these post-transitional modifications, increased ribosomal biogenesis can occur as evidenced by assessment of nucleolar size, qRT-PCR, and proliferation and growth of hepatocellular carcinoma (Zhang et al., 2023; Sun et al., 2024). These findings link *fasn-1* to enhanced rRNA synthesis through malonyl-cOA in cancer models, a key metabolite involved in the regulation of fatty acid metabolism as well as nutrient sensing.

Investigation into how biguanides regulate rRNA synthesis could be important to our understanding of the safety and efficacy of these drugs as potential anti-aging interventions. Transcription of rRNA is negatively regulated by nucleolin (*ncl-1*) and thus *ncl-1* null mutants exhibit increased nucleolar size, higher levels of rRNA, and translate more protein (Frank & Roth, 1998). Unleashing rRNA synthesis by knocking down *ncl-1* blocks several lifespan extending paradigms associated with reduced TORC activity, insulin signaling, reproduction, and mitochondrial activity in *C. elegans* (Tiku et al., 2017). Metformin-lifespan extension has been shown to act through a number of these distinct gero-protective pathways (Soukas, Hao & Wu, 2019), thus it seems logical that reduced rRNA synthesis would be necessary for phenformin-mediated lifespan extension.

We demonstrate that increasing rRNA synthesis results in heightened sensitivity to the effects of combined phenformin and *fasn-1* deficiency. Notably, the stress reporter for *nlp-*29 was activated in *ncl-1* deficient animals upon treatment with phenformin with or without genetic inhibition of *fasn-1*. Combining all three conditions resulted in synergistic overexpression of *nlp-29* supporting our hypothesis that enhanced rRNA synthesis hypersensitizes animals to the effects of phenformin-induced reductive stress. Together, these findings suggest that reductions in nucleolar activity may be necessary for the detoxification of the drug, highlight the regulatory role of the nucleolus in aging, and point to nucleolar activity being the biological switch that determines whether phenformin is a life-extending or life-shortening intervention.

It remains unclear whether enhanced rRNA synthesis is a cause or consequence of reductive stress. Hypoxia has been shown to cause the translocation of hypoxia inducible factor *HIF1a* into the nucleolus where it promotes rRNA transcription, driving cancer growth and proliferation in a variety of cellular breast cancer lines (Elhamamsy et al., 2025). Hypoxia is known to cause reductive stress in some instances due to a shutdown of oxidative phosphorylation (Ravi, 2025). Thus, hypoxia is an example of a situation where reductive stress could drive rRNA synthesis.

If enhanced rRNA synthesis does cause an increase in basal antioxidant levels, it could be a pathway through which oxidative stress could be addressed. Intriguingly, previous work finds that increased ribosome biogenesis occurs in the blood leukocytes of patients with schizophrenia as evidenced by increased rDNA copy number and rRNA transcripts (Ershova et al., 2025). Ribosome biogenesis was found to be negatively correlated with markers for oxidative stress induced cellular damage, as suggested by longer telomere length, fewer DNA breaks, and reduced oxidation of DNA (Ershova et al., 2025). This was attributed to a negative correlation between rRNA biogenesis and the pro-oxidant enzyme NOX4, suggesting that enhanced rRNA biogenesis may be an adaptive response against oxidative stress in the blood cells of schizophrenic patients (Ershova et al., 2025). This seems to suggest that increasing ribosome biogenesis could shift redox ratios towards a reduced state by inhibiting oxidant-producing pathways. Future work should be done to determine whether increasing rRNA synthesis results in an accumulation of antioxidants and whether this can improve resistance to oxidative stressors. Such studies could provide new therapeutic insight into the possibility of manipulating redox homeostasis through a highly conserved rRNA synthesis pathway.

Existing literature has already identified differential rRNA modification as a key regulator of various stress responses and longevity. For example, preventing the methylation of 18S rRNA by *dimt-1* at adenosines 1735 and 1736 in *C. elegans* is sufficient to not only extend lifespan up to 33%, but can also improve resistance to UV irradiation, and heat stress via selective binding of genes involved in longevity, degradation pathways, detoxification, glutathione metabolism and energy production (Rothi et al., 2025). Similarly, mutation of *metl-5* prevents the methylation of adenosine 1717 on 18S rRNA, resulting in increased resistance to oxidative, cold, severe heat shock, and UV irradiation thanks to the reduced translation of the gene *cyp-29A3* (Liberman et al., 2020). These findings highlight the importance of ribosome diversity in regulating cellular responses to stressors and other stimuli. Variation in rRNA processing is a major source of ribosome diversity and plasticity that is often underappreciated. Thus, ribosome heterogeneity provides an added level of post-transcriptional gene regulation by enabling selective loading and translation of mRNA transcripts (Trahan & Oeffinger, 2023).

Quality control of RNA in the nucleus and cytoplasm involves the RNA exosome complex consisting of 9 subunits (exosc-1-9) that make up the exosome’s inert core and two catalytic subunits (*exosc-10* and *DIS3*) play a dual role in the degradation of erroneous mRNA and rRNA transcripts as well as the processing of rRNA (Puno et al., 2019). Knockdown of EXOSC2 and EXOSC5 through EXOSC10 resulted in impaired processing of 5.8S and 18S rRNA but not other RNA exosome subunits or coactivators (Tafforeau et al., 2013; Feng et al., 2025). Thus, targeting specific subunits of the RNA exosome may also be a source of ribosome diversity and a way to boost an organism’s resistance to stress.

Our initial reverse genetic screen revealed that RNAi knockdown of either *crn-3* or *bud-23* was sufficient to significantly reverse the reductive death associated with combined *fasn-1* deficiency and phenformin treatment. Specifically, *crn-3* is part of the RNA exosome which is involved in the processing of the 5.8S and 18S rRNA (Tafforeau et al., 2013; Zhu et al., 2018). Meanwhile, *bud-23* is a rRNA methyltransferase which aids in the processing of 18s rRNA (Zhu et al., 2018). Further exploration revealed that knocking down genes involved in the nucleolar transcription, processing in the nucleoplasm, nuclear exportation, and cytoplasmic processing had the same protective effect against reductive stress. Thus, our findings support existing work showing that rRNA modification is a key pathway through which stress can be mitigated. Interestingly, we find that directly inhibiting rRNA transcription through *tif-1a* also improved stress resistance. Aberrant rRNA synthesis caused by the targeted knockdown of the RNA exosome and rRNA modifying or processing can cause a decrease in overall rRNA levels by inhibiting RNA polymerase I activity via a nuclear RNAi mediated pathway that targets rRNA (Zhu et al., 2018, Liao et al., 2021). Thus, it’s possible that the aberrant modification of a single rRNA nucleotide can cause a decrease in the overall abundance of rRNA, protecting against reductive stress.

Notably, we found that *rsks-1* was necessary for rRNA-mediated reductive stress resistance. These findings were surprising as other investigators have suggested a feedback loop in which mutating TOR-pathway associated genes *rsks-1* or *rict-1* prevents the nucleolar localization of some RNA exosome subunits and vice versa where inhibition of RNA exosome subunits downregulate translation via reduced TOR - signaling (Feng et al., 2025). Others have asserted that boosting mTOR activity promotes *tif-1a* association with rDNA, increasing rRNA synthesis (Mayer et al., 2004). As such, we would have expected a non-additive effect when treating *rsks-1* mutants with either *crn-3* or *tif-1a* RNAi.

We wonder whether the benefits of inhibition of rRNA biosynthesis are dependent upon the selective translation of mRNA transcripts. Previous work has shown that inhibition of 18s methylation by *metl-5* is sufficient to dramatically alter the polysome profiles, promoting the association of mRNA transcripts with ribosomes that encode proteins that mobilize cellular defenses against stress (Liberman et al, 2020). As such, attenuation of translation via the administration of cycloheximide eliminated the stress resistance observed in *metl-5* deficient *C. elegans* (Liberman et al, 2020). These findings seem consistent with our own observations, leading us to speculate that *crn-3* mutation could trigger selective loading of reductive stress responsive - mRNAs onto polysomes, thus requiring sustained activity of *rsks-1*.

Previous work has shown that inhibiting nucleolar activity can improve resistance by changing the lipidome (Sharifi et al., 2024). Inducing nucleolar stress can activate transcription factor *pha-4* and the resulting increase in stored fat improves starvation resistance in *C. elegans* (Feng et al., 2025; Wu et al., 2018). Similarly, inhibition of tRNA genes which are essential for ribosomal translation can also promote lipid accumulation and starvation resistance through an AMPK signaling pathway (Webster et al., 2017). The type of fatty acids that is produced may also dictate resistance to specific stressors. For example, impairing *crn-3* in *C. elegans* enhances production of oleic acid to improve oxidative stress resistance (Feng et al., 2025).

Consistent with previous work, we observed that *crn-3* deficient animals overexpress FASN-1 protein. In the context of reductive stress, enhanced *fasn-1* activity associated *crn-3* deficiency is likely to be advantageous as *de novo* fatty acid synthesis is an essential pathway through which NADPH is consumed. Thus, *crn-3* deficiency and reduced rRNA synthesis may beneficially increase redox buffering capacity. However, the current study focuses on how nucleolar stress can improve stress resistance in the absence of *de novo* fatty acid synthesis activity due to the utilization of RNAi and auxin-inducible-degron (AID) knockdown systems. We show that *crn-3* deficient animals were resistant to reductive stress even when FASN-1 protein was depleted. Intriguingly, our findings suggest that knockdown of rRNA processing genes can significantly buffer against reductive stress in a manner that is independent of their regulatory roles over lipid metabolism.

## Limitations

Future work should also be done to determine if alterations to rRNA biogenesis can rescue toxicity associated with other paradigms of reductive stress and not just phenformin-induced reductive stress. Additionally, this study focuses on inhibiting ribosomal biogenesis by targeting of RNA polymerase I activity and RNA polymerase I transcribed rRNAs. It would be interesting to determine whether interrupting ribosome biogenesis via the perturbation of RNA polymerase III-mediated 5S rRNA and tRNA synthesis or ribosomal proteins may similarly protect against reductive stress.

## Conclusions

Overall, our findings suggest that restricting pre-rRNA transcription or altering its modification and processing may serve to detoxify reductive stress. Targeting this highly conserved pathway may beneficially increase buffering capacity during insults to redox homeostasis. As such, rRNA biogenesis could hold therapeutic potential for addressing diseases such as cancers, cardiovascular disease, and other metabolic ailments associated with reductive stress.

## Methods

### Nematode Strains and Maintenance

*C. elegans* strains were maintained by feeding OP50-1 *ad libidum* on Nematode Growth Medium (NGM) agar plates at 20°C in a temperature and humidity controlled Thermo Scientific Forma incubator, unless otherwise noted (Brenner, 1974). Ancestral vials for each strain were freshly thawed every 3-4 months to prevent genetic drift in existing cultures. The strains used are listed below

| Strain | Genotype | Notes |
| --- | --- | --- |
| N2 |  | Ordered from CGC |
| MGH770 | *osIs158[eft-3p::AtTIR1(F79G)::mRuby] II; [GFP::AID\*::3x-FLAG::fasn-1]* | Constructed for this paper |
| MGH887 | *crn-3(ok2269)* | VC1715 ordered from CGC and backcrossed 4x to N2 |
| MGH888 | *crn-3(ok2269); osIs158[eft-3p::AtTIR1(F79G)::mRuby] II; [GFP::AID\*::3x-FLAG::fasn-1]* | Constructed for this paper. MGH887 crossed into MGH770 |
| MGH9 | *rsks-1(ok1125)* | RB1206 ordered from CGC and backcrossed 8x to N2 |
| MGH989 | *rsks-1(ok1255); [GFP::AID\*::3x-FLAG::fasn-1]; osIs158[eft-3p::AtTIR1(F79G)::mRuby] II* | Constructed for this Paper. MGH9 backcrossed into MGH770 |
| IG274 | *frIs7 [nlp-29p::GFP + col-12p::DsRed] IV* | Ordered from CGC |
| MGH823 | *fasn-1(fr8) I; frIs7 [nlp-29p::GFP + col-12p::DsRed] IV* | IG348 Ordered from CGC and backcrossed 8x to N2 |
| *bud-23(5768)* | *C27F2.4/bud-23(5768)* | Gifted by Greer Lab,<br>Washington University St.<br>Louis |
| COP262 | *unc-119(ed3)*<br>*III; knuSi221 [fib-1p::fib-1(genomic)::eGFP::fib-1 3' UTR]* | Ordered from CGC |
| QK52<br>(previously<br>JM43) | *rde-1(ne219)*<br>*V; xkIs99 [wrt-2p::rde-1::unc-54 3'UTR]* | Gifted by Ruvkun Lab,<br>MGH |
| MGH167 | *sid-1(qt9); alxIs6[VHA-6p::SID-1::SL2::GFP]* |  |

### Alkaline Hypochlorite Synchronization of *C. elegans*

To synchronize worms for experiments, gravid animals which were collected in M9 medium and centrifuged at 4400 g for 30 seconds and resuspended in 6 mL of 1.3% bleach, 250 mM NaOH for 1 minute and shaken vigorously. The first bleach exposure was followed by centrifugation for 30 seconds and a wash in M9. A second bleach exposure was performed followed by 3-4 more washes in M9 to remove residual bleach, Eggs were resuspended in 7-12 mL of M9 and left to rotate for 16-24 hours at 20°C, allowing synchronized L1 larvae to hatch.

### *E. coli* Strains and Maintenance

The *E.* coli strain used for general worm maintenance and non-RNAi experiments was OP50-1. Cultures of OP50-1 were grown in LB broth for 15 - 24 hours at 37°C without shaking and were seeded directly onto NGM plates. In general, plates were used 3-7 days after seeding. All RNAi experiments were conducted using *E. coli* HT115 (DE3) as a vector. Cultures were grown in LB broth for 15-24 hours at 37 °C with shaking. Cultures were pelleted via centrifugation and excess fluid was aspirated in order to achieve a 5x concentration. Bacterial cultures were seeded directly onto NGM plates containing 200 µg/mL carbenicillin and 5 mM isopropyl-B-D-thiogalactopyranoside (IPTG). Plates were seeded at least 1 day before they were used to ensure sufficient IPTG induction and used within 5 days of seeding.

### Lifespan Analysis

Strains were grown on OP50-1 until gravid and then washed off the plates in M9 buffer. Worms were synchronized via alkaline hypochlorite treatment and left on a rotator at 20 °C for 18-24 hours. L1 worms were dropped onto OP50-1 plates and allowed to grow to the L4 stage. A total of 50 - 60 L4 worms were transferred to each experimental plate which was treated with 100 µM 5-fluorodeoxyuridine (FUdR) to suppress progeny development. The total number of worms transferred to each plate was between 50-60. All lifespans were conducted at 20°C and were post-developmentally treated with RNAi and drugs unless otherwise specified.

### Development Analysis

RNAi plates were prepared as described above and drug-treated over the top of the bacterial lawn and dried in a laminar flow hood. The appropriate strains were grown and synchronized to the L1 stage as described above before being dropped directly onto the appropriate experimental plates. Worms were grown at 20 °C. At the indicated time points, animals were scored for their transition into adulthood by the appearance of the vulva slit.

### 96-well Metabolism RNAi Library Screen

96-well RNAi high throughput microscopic assessment of animals was performed as previously described (Pino et al., 2013). Larval development of eft-3p::AtTIR1(F79G)::mRuby; 3xFLAG::AID*::GFP::FASN-1 animals was assessed following at least 72 hours of combined 4.5 mM phenformin and 5 µM 5-PH-IAA treatment. Firstly, animals were bleach synchronized for 18 hours, and L1s were subsequently dropped onto 96-well RNAi bacto agar plates pretreated with 4.5 mM phenformin and 5 µM 5-PH-IAA (to induce degradation of FASN-1) and pre-seeded with 1,046 individual RNAi clones (corresponding to genes annotated as metabolism-related according to Gene Ontology terms). Animals were then incubated for 72 hours at 20°C prior to microscopic analysis. The number of viable adults were counted per well under dissecting microscope and compared to the percent reaching adulthood with empty vector (EV) control. The top 91 RNAi clones with the greatest number of viable adults were selected for re-testing on larger 60 mm x15 mm petri dishes with an additional series of EV control plates to identify candidate targets. Visualization of the screen results were performed using GraphPad Prism 10.

### qRT-PCR Analyses

qRT-PCR was performed as previously described (Cedillo et al., 2023) Approximately 1000 synchronized animals of the stage and treatment conditions indicated in the figure legends were harvested using M9 buffer and washed an additional 3x times to remove residual feeding bacteria. Animals were homogenized using a Qiagen TissueLyzer II with metal beads, centrifuged for 10 minutes at 12,000xg to pellet cuticle and cellular debris, and RNA was extracted from the supernatant using RNAzol RT (Molecular Research Center) according to manufacturer instructions. RNA was treated with RNase free DNAse prior to reverse transcription with the Quantititect Reverse Transcription kit (Qiagen). qRT-PCR was conducted using a Quantitect SYBR Green PCR reagent (Qiagen) following manufacturer instructions on a Bio-Rad CFX96 Real-Time PCR system (Bio-Rad). Expression levels of tested genes were presented as normalized fold changes to the abundance control genes (as indicated in the Figure Legends) using the ΔΔCt method (Livak & Schmittgen, 2001). The following primers were used for expression analysis:

*bud-23*, Forward: 5’ ATGAAATGGCTGAAAGAGCAC’

*bud-23*, Reverse: 5’ TACCGCAACCAATATCTAGCA 3’

*crn-3*, Forward: 5’ GAATGCACTCATGAACAAAGTC 3’

*crn-3*, Reverse: 5’ TAATGTTCGACTGATGAACCG 3’

eft-3. Forward: 5’ GTAAGGGATCTTTCAAGTACGC 3’

eft-3, Reverse: 5’ CATCGATGATGGTGATGTAGTAC 3’

### Puromycin Incorporation Assay

Puromycin incorporation was performed in *C. elegans* as previously described with some modifications (Clay et al., 2023). Overnight cultures of the appropriate RNAi clones were induced with IPTG for 2 hours at a concentration of 175 µL 1 M IPTG per 35 mL of bacterial culture. Bacterial cultures were then centrifuged and resuspended at a 10x concentration. Worms were incubated in a 1.5 microcentrifuge tube with 200 µL of 10x bacteria, 790 mL of S-buffer, and 10 µL of 50 mg/mL puromycin (final concentration 0.5 mg/mL) for 4 hours at 20°C on a rotator or rocker. Worms were then rinsed 3 times in PBS and flash frozen in liquid nitrogen and stored hours at −80°C prior to processing for immunoblotting.

### Immunoblotting

Worm and cell lysates were prepared in RIPA buffer supplemented with protease and phosphatase inhibitors (Roche cOmplete™, EDTA-free Protease Inhibitor Cocktail, Sodium pyrophosphate, sodium fluoride, sodium orthovanadate, β-Glycerophosphate, sodium Molybdate, and Okadaic acid). Samples were sonicated at 4°C using a Q-sonica set at 40% amplitude at intervals of 30s/30x on and off for a total of 20 minutes. Samples were then spun down for 15 minutes at 14,000 rpm at 4°C to pellet debris and the supernatant was transferred to a fresh tube. A Bicinchoninic Acid (BCA) assay was used to quantify the protein in the supernatant of each sample. The concentration of the samples was then normalized to the sample with the lowest concentration, using RIPA lysis buffer as a dilutant. 4x Laemmli sample buffer (Bio Rad) supplemented with 10% (v/v) β-mercaptoethanol was added to each sample for a final 1x concentration. Samples were boiled at 95°C for 5 minutes and then stored at 20°C until used for SDS-PAGE. Protein samples were loaded into 4-15% Mini-PROTEAN® TGX™ Precast Protein Gels (Bio Rad) and run at 150 volts for 1-1.5 hours in a Bio-Rad Mini Protean Tetra Cell gel system. Bio-Rad’s Precision Plus Dual Color standard was used as a protein ladder. Proteins were transferred to a nitrocellulose membrane via electrophoretic transfer using a wet tank-transfer system run at 100V for 1 hour at 4°C in a 20% methanol transfer buffer. Ponceau staining was used to visualize proteins before blocking for 1 hour in 5% milk dissolved in TBST. Primary antibody incubation was conducted overnight at 4°C. The following antibodies were used: mouse monoclonal anti-Actin (C4) (Abcam, Cat# ab14128, 1:2500), rabbit monoclonal anti-GFP (D5.1) (Cell Signaling Technology, Cat# 2956, 1:1000), mouse monoclonal anti-puromycin (12D10))(Millipore Sigma, Cat# MABE343, 1:1000). Membranes were then washed 3 times with 1x TBST. For primary antibody immunodetection, a 1:5000 dilution of Goat anti-rabbit HRP conjugate or Goat anti-mouse HRP conjugate (GE Heathcare) in a 5% bovine serum albumin (BSA) 1x TBST solution was used, incubating membranes for 1 hour rocking at room temperature prior to the addition of SuperSignal West-Pico chemiluminescent substrate (Thermo Pierce). Blots were visualized using an Invitrogen iBright Imaging system. To reprobe blots for the loading control, 10 mL of thermoFisher’s Restore™ PLUS Western Blot Stripping Buffer was added to cover the membrane and left to rock at room temperature for 10 minutes before washing once in TBS and then re-blocking in 5% milk in TBST for 1 hour. Blots could then be incubated in the desired primary antibody overnight as previously described. Western blots were quantified for band intensity using ImageJ software’s rectangle selection tool. Significance between different treatment groups was determined using a one-way ANOVA, accounting for multiple comparisons via Sidak corrections using Graphpad’s Prism 10 software. Comparisons were considered significant if p < 0.05.

### Live Worm Imaging/Quantification of Fluorescence Intensity

Worms were mounted onto 2% agarose pad and sedated with 2.5 mM levamisole and imaged using a Leica DM6 B microscope with Thunder Imaging capabilities under the appropriate brightfield or fluorescent channel and magnification. Images were quantified by manually tracing animals using ImageJ software’s polygon selections tool. Significance was determined using a one-way ANOVA, accounting for multiple comparisons using Sidak corrections using Graphpad’s Prism 10 software. Comparisons were considered significant if p < 0.05.

### Fixed Nucleolar Imaging in C. elegans

Staining for the imaging of nuclei and nucleoli was performed as previously described in Sharifi et al. (2024) with a few modifications. Worms were collected by washing plates and rinsed 3 times to remove residual bacteria. In a 1.5 µL microcentrifuge tube, worms were fixed in 1 mL of 50% ethanol (diluted in water) for 5 minutes and then washed in PBS three times. Worms were then incubated in 1 mL of 10 µM hoechst 3342 (final concentration 0.5 ul/mL) for 15 minutes in the dark. Worms were mounted directly onto glass microscope slides and imaged at 63x magnification under oil immersion using a Leica DM6 B microscope. The nuclear stain hoechst 3342 was imaged under a BFP channel while the nucleolar *fib-1:GFP* signal was imaged under a GFP channel. Images were quantified by manually tracing the nuclei and nucleoli using ImageJ software’s polygon selections tool. At least 3 hypodermal cells per worm were imaged. Nucleolar size was calculated by dividing the nucleolar area of each cell by its nuclear area, thus normalizing to each individual cell. The average nucleolar to nuclear ratio for each worm was then calculated. When comparing different conditions, significance was determined using a one-way ANOVA, accounting for multiple comparisons using Sidak corrections on Graphpad’s Prism 10 software. Comparisons were considered significant if p < 0.05.

### RNA-sequencing analyses

For bulk RNA-sequencing analyses, 1000 worms of the indicated strain, genotype, or treatment per sample and replicate was harvested using M9, washed three times to remove bacteria, and worm pellets were resuspended into 500 microliters of RNAzol RT. Total RNA was subsequently extracted and the genomic DNA was removed using the Direct-Zol RNA Miniprep Plus Kit (Zymo Research) following manufacturer’s instructions. Total extracted RNA was evaluated for quality control using a NanoDrop One C Microvolume UV-Vis Spectrophotometer (Thermo Fisher). Samples were sent to Azenta (Genewiz) for additional quality control, library preparation, and mRNA sequencing. Samples were validated for RNA integrity with a RIN score > 9 and DV200 > 70 using an RNA Tapestation 4200 (Agilent). Illumina library preparation was performed using polyA selection for mRNA species. Approximately 20 million paired-end 150bp reads were generated per sample, with ≥ 80% of bases passing a Phred quality score ≥ Q30. Fastq read quality control, adapter trimming, quality score filtering, and quasi-alignment were all performed using custom UNIX/Bash shell scripts on the Mass General Brigham ERISTwo Scientific Computing CentOS 7 Linux Cluster. Reads were analyzed for quality control using FastQC v0.11.8 (http://www.bioinformatics.babraham.ac.uk/projects/fastqc) and MultiQC v1.19^81,82^. Reads were then filtered for Illumina adapter contamination, truncated short reads or low-quality base calls using BBDuk (BBTools). The subsequently trimmed and cleaned reads were then quasi-aligned to the Caenorhabditis elegans reference transcriptome annotation (WBcel235, Ensembl Release 105) using Salmon v1.9.0, correcting for GC content and sequencing bias using the command parameters ‘--gcBias’ and ‘--seqBias’ (Patro et al., 2017). All statistical analyses and visualizations were generated using the R v4.3.2 (www.r-project.org) Bioconductor v3.18 statistical environment on a local machine through Jupyter Notebook v6.4.10 (https://jupyter.org) (Huber et al., 2015). Quasi-aligned transcript quantification files for each sample were collapsed into gene-level count matrices using R package tximport v1.30.0 (Soneson, Love & Robinson, 2015). Paired differential expression was calculated using R package DESeq2 v1.42.1 with a design formula of ‘∼ RNAi_Treatment’ (Love, Huber & Anders, 2014). Genes were considered differentially expressed with a Benjamini-Hochberg False Discovery Rate (FDR) corrected P value < 0.05 and an absolute log2 transformed fold change of 1 (Benjamini & Hochberg, 1995). Comparison of differentially expressed genes with prior *C. elegans* mutant datasets was performed using WormExp v2.0 (Yang, Dierking & Schulenburg, 2016). Tissue expression prediction analysis of differentially expressed genes as indicated in the figure legends was performed using the Worm Tissue Prediction Server and data plotted using Graphpad Prism 10 (Kaletsky et al., 2018). Visualizations were post-edited for font, sizing, and appearance using Adobe Illustrator and the Adobe Creative Cloud Suite. All RNA-sequencing data has been uploaded to the NCBI Gene Expression Omnibus (GEO) under accession number GSE277946.

### NADPH/NADP and NADH/NAD+ Ratiometric Profiling

NADPH/NADP and NADH/NAD+ ratiometric profiling: Ratiometric redox profiling was performed using the NADP/NADPH Ratio Red Fluorescence Assay Kit (eEnzyme, Cat# CA-N426) and the NAD/NADH Colorimetric Assay Kit (abcam, Cat# ab65348) according to manufacturer’s instructions. To prepare lysates for reducing equivalent extraction, ∼1000 nematode lysates for each condition and sample indicated were collected into M9 buffer supplemented with EDTA-free protease inhibitor tablets (Roche) and a phosphatase inhibitor cocktail containing sodium fluoride, sodium pyrophosphate, sodium orthovanadate, β- glycerophosphate, and okadaic acid. Samples were then sonicated at 40% amplitude in a QSonica Q800R3 water bath sonicator at 30 s ON/30 s OFF intervals for a total of 20 minutes at 4°C, ensuring cuticle breakdown uniformly across each sample via dissecting microscope. Lysates were cleared by centrifugation at 21,000xg at 4°C for 10 minutes and supernatants were retained in clean microcentrifuge tubes for NADPH/NADP+/NADH/NAD+ extraction. Fluorescence emission spectra were collected using a Spectramax i3x plate reader, and colorimetric measurements were collected using a Spectramax PLUS plate reader. Values were plotted and significance was determined using Graphpad Prism 10.

### Statistical analysis

To determine statistical significance for developmental assays and quantification of fluorescence or western blot band intensity, GraphPad’s Prism 10 software was used to run one-way and two-way ANOVAs. Unless otherwise noted, all ANOVAs were run with sidak correction for multiple comparison testing and any comparisons shown in figures report the corrected p-values where *p < 0.05* was the threshold for significance. Asterisks denote corresponding statistical significance ∗p < 0.05; ∗∗p < 0.01; ∗∗∗p < 0.001 and ∗∗∗∗p < 0.0001.

For lifespan analysis experiments, the software OASIS 2 was used to determine statistical significance between the mean lifespans of different experimental conditions using Kaplan-Meier estimation and log-rank testing. Unless otherwise specified, comparisons shown in figures and tables report the Bonferroni corrected p-value where values where *p < 0.05* was the threshold for significance. Asterisks denote corresponding statistical significance ∗p < 0.05; ∗∗p < 0.01; ∗∗∗p < 0.001 and ∗∗∗∗p < 0.0001.

## Data availability statement

Any strains or plasmids used in this study are available upon request. All data necessary for confirming the conclusion of the article can be located within the article, figures, and tables. All RNA-sequencing data has been uploaded to the NCBI Gene Expression Omnibus (GEO) under accession number GSE277946.

## Acknowledgements

We thank Dr. Eric Greer (Washington University in St. Louis) and the Caenorhaditis Genetic Center (CGC) (Univeristy of Minnosota) for providing us with strains for this study. The CGC is funded by the NIH Office of Research Infrastructure Programs (P40 OD010440). This work was funded by the NIH (R01 AG058259 and R01 AG69677 to A.A.S.), the Nutrition Obesity Research Center at Harvard (NORCH) (P30 DK040561 to A.A.S.), the NIH/NIDDK-funded Boston-Area DERC (P30 DK135043 to A.A.S.), Funding from the National Institute on Aging also supported this work (K08 AG078447 to A.Y.)

## Author Contributions

J.R., conceptualization, investigation, formal analysis, methodology, validation, and writing-original draft; F.A., conceptualization, investigation, formal analysis, methodology, review, and editing; N.L.S., review and editing; S. E., review and editing; A.S., supervision, funding acquisition, review, and editing; A.Y., supervision, funding acquisition, review, and editing.

