## Supplemental Data for "Ribosomal RNA Synthesis is a Lethal Vulnerability During Reductive Stress In *C*. *elegans*"

### Membrane 1

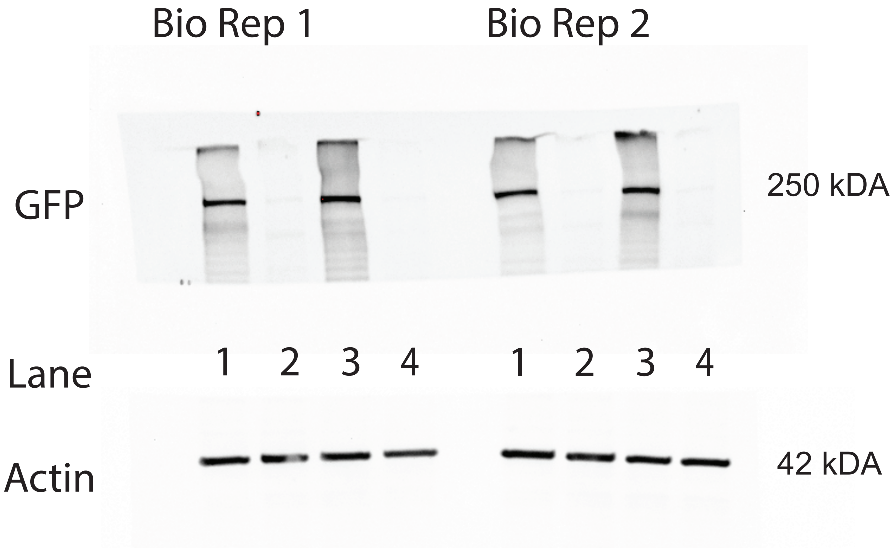


### Membrane 2

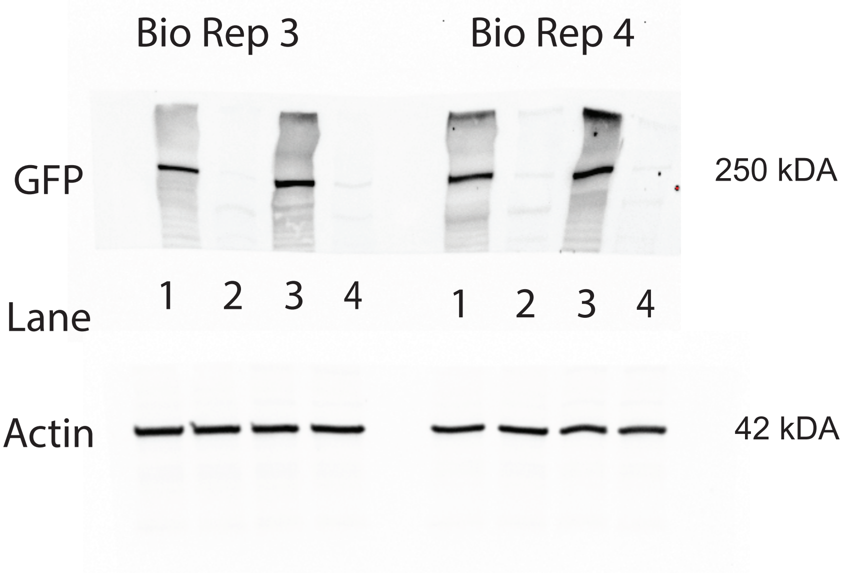


Lane/Strain+Treatment ***All samples collected at AD2

*1. eft-3p::AtTIR1(F79G)::mRuby; 3xFLAG::AID*::GFP::FASN-1*, OP50-1, vehicle, 0.1% DMSO

2. *eft-3p::AtTIR1(F79G)::mRuby; 3xFLAG::AID*::GFP::FASN-1*, OP50-1, vehicle, 5 μM 5-Ph-IAA

*3. crn-3(ok2269); eft-3p::AtTIR1(F79G)::mRuby; 3xFLAG::AID*::GFP::FASN-1*, OP50-1, vehicle, 0.1% DMSO

4. *crn-3(ok2269);eft-3p::AtTIR1(F79G)::mRuby; 3xFLAG::AID*::GFP::FASN-1*, OP50-1, vehicle, 5 μM 5-Ph-IAA

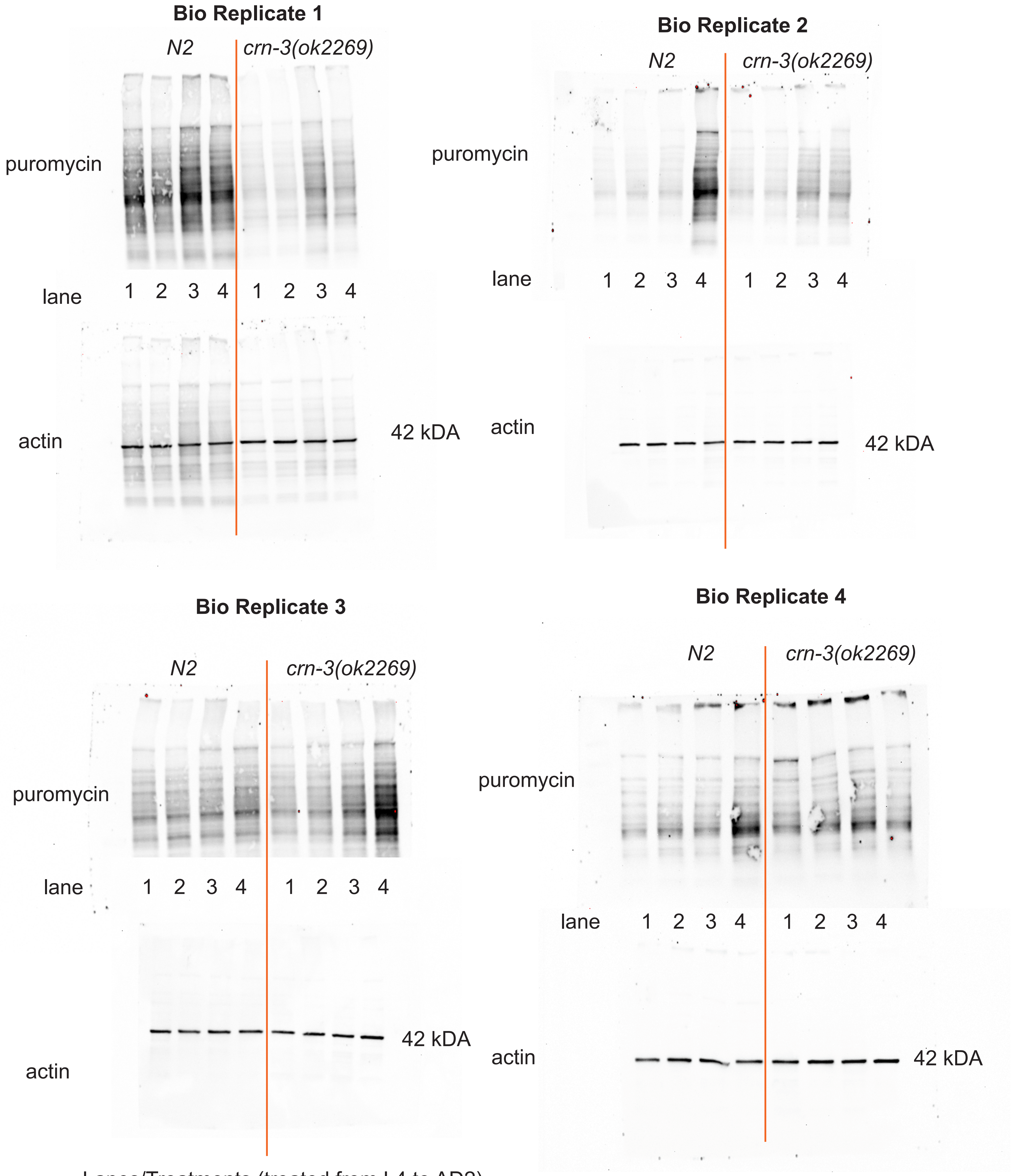


Lanes/Treatments (treated from L4 to AD2)

1. EV, Vehicle
2. EV, 4.5 mM Phenformin
3. *fasn-1* RNAi, Vehicle
4. *fasn-1* RNAi, 4.5 mM Phenformin
