## Supplemental Figures S1-S4 for "Ribosomal RNA Synthesis is a Lethal Vulnerability During Reductive Stress In *C*. *elegans*"

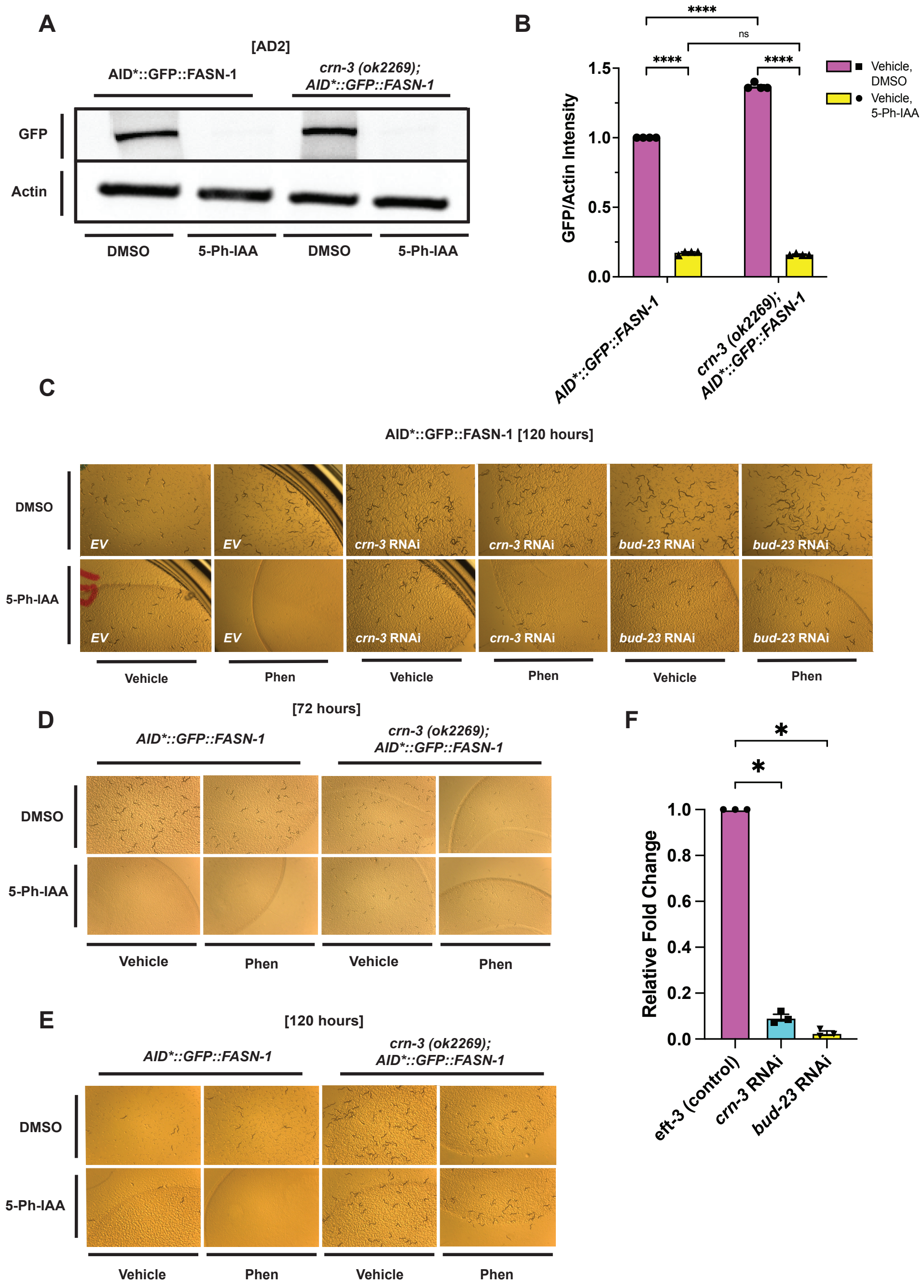


**Supplementary Figure 1. Knockdown of *crn-3* and *bud-3* via Auxin Degradable Degron System or RNA Interference Protects Against Phenformin-Induced Reductive Stress in *Fasn-1* Depleted Animals**

(A) Western blot showing abundance of FASN-1 protein in *eft-3p::AtTIR-1(F79G)::mRuby;3xFLAG::AID*::GFP::FASN-1 and crn-3(ok2269)*; *eft-3p::AtTIR-1(F79G)::mRuby;3xFLAG::AID*::GFP::FASN-1* upon treatment with either water vehicle control or 4.5 mM phenformin with or without 0.1% or 5 µM 5-Ph-IAA. Results are representative of 4 independent biological replicates. Note that *eft-3p::AtTIR-1(F79G)::mRuby;3xFLAG::AID*::GFP::FASN-1* were treated with the respective drugs from the L4 stage to AD2 due to 5-Ph-IAA treatment being toxic from L1 of hatching. Meanwhile *crn-3(ok2269)*;*eft-3p::AtTIR-1(F79G)::mRuby;3xFLAG::AID*::GFP::FASN-1* animals were treated with the respective drugs from L1 of hatching to AD2.

(B) Quantification of FASN-1 protein expression in control and *crn*-3 mutants

(C) Representative images of developmental assay performed on *eft-3p::AtTIR-1(F79G)::mRuby;3xFLAG::AID*::GFP::FASN-1* animals treated with a water vehicle or 4.5 mM phenformin with or without 0.1% dmso or 5 μM 5-Ph-IAA.

(D - E) Representative images of developmental assay performed on control *eft-3p::AtTIR-1(F79G)::mRuby;3xFLAG::AID*::GFP::FASN-1* or *crn-3(2269); eft-3p::AtTIR-1(F79G)::mRuby;3xFLAG::AID*::GFP::FASN-1* animals after (D) 72 hours of treatment or (E)120 hours of treatment from L1 of hatching.

(F) Quantification of RT-qPCR measuring RNAi knockdown efficiency of *crn-3* and *bud-23* mRNA transcripts compared and normalized to an *eft-3* control. Results are compiled from 3 independent biological replicates. All error bars represent mean +/- SEM unless otherwise noted. Significance was assessed in (B-C) using a two-way ANOVA with Sidak correction of multiple comparison testing. ns $p > 0.05$, * $p < 0.05$, ** $p < 0.01$, *** $p < 0.001$, **** $p < 0.0001$.

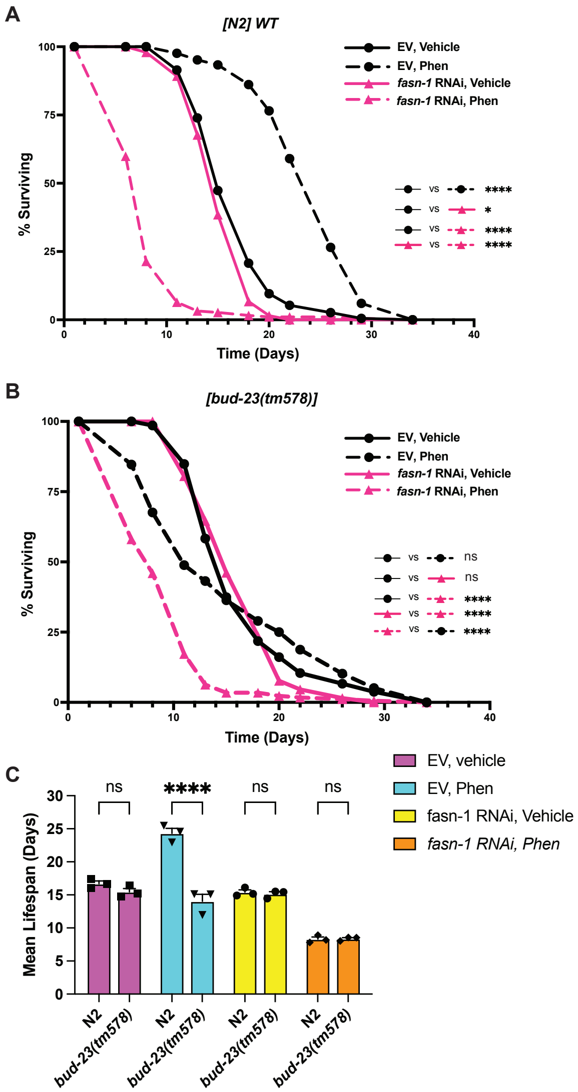

### Supplementary Figure 2. Genetic Mutation of *bud-23* Does not Reverse Post-Developmental Reductive Death

(A-B) Lifespan analyses of (A) wild-type and (B) *bud-23(tm578)* animals.
(C) Mean lifespans of N2 wild-type animals vs *bud-23(tm578)* animals. Data represents 3 independent biological replicates n = 3. All mutant animals were backcrossed 4-6 times to wild-type N2 animals to remove potential background mutations. Significance was assessed in (A-C) using log rank analyses. ns p > 0.05, * p < 0.05, ** p < 0.01, *** p < 0.001, **** p < 0.0001. See Table S2 for log rank analyses.

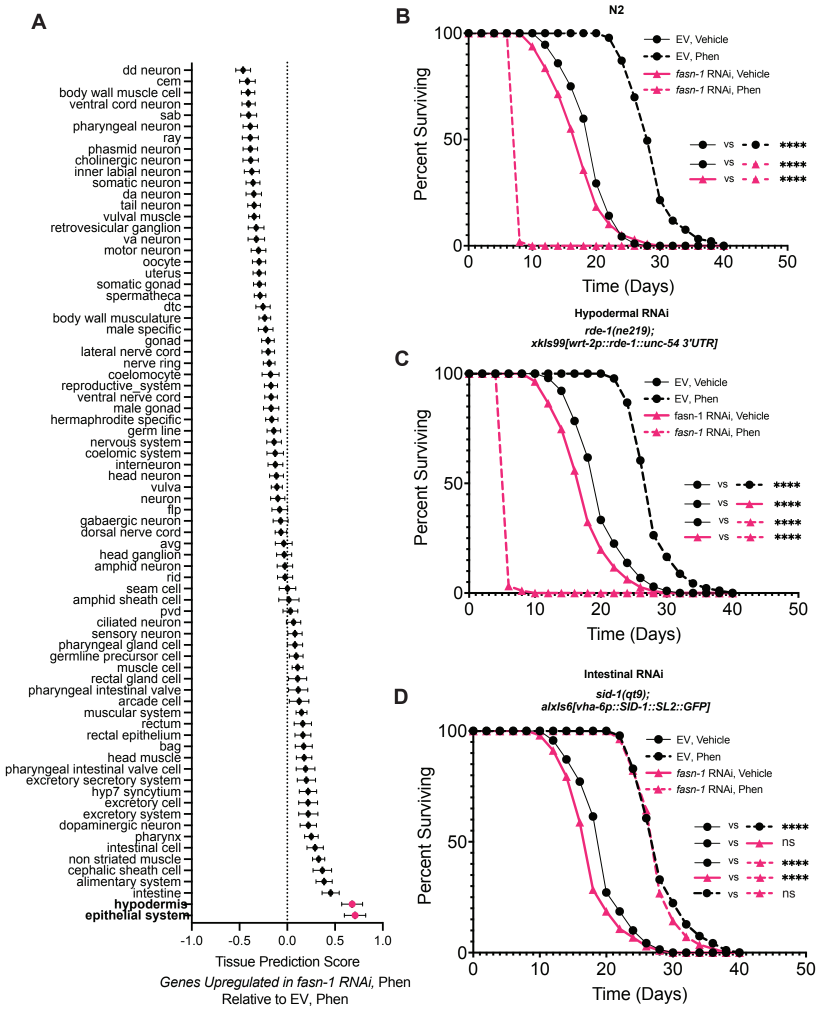


### Supplementary Figure 3. Hypodermal Expression of *fasn*-1 Protects Against Reductive Death

(A) Tissue prediction analysis of genes differentially upregulated by *fasn-1* RNAi with 4.5 mM phenformin relative to EV with 4.5 mM phenformin. Genes were selected based on a $\log_2$ fold change greater than 1 and an adjusted *p* value < 0.05. Tissue prediction scores were generated for the selected list using https://worm.princeton.edu.

(B-D) Lifespan analysis of (B) N2 wildtype C) hypodermal specific RNAi competent animals (D) intestinal specific RNAi competent animals treated with or without *fasn-1* RNAi and 4.5 mM phenformin from L4 to end of life. Significance was assessed for **(A)** using DESeq2 Wald testing with Benjamini-Hochberg FDR correction, and (B-D) using log rank analyses as shown in Table S3. ns $p > 0.05$, * $p < 0.05$, ** $p < 0.01$, *** $p < 0.001$, **** $p < 0.0001$.

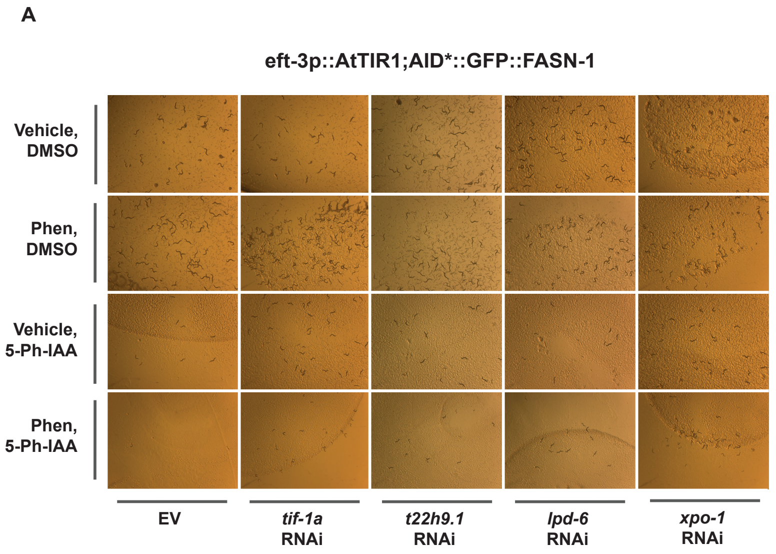


### Supplementary Figure 4. RNAi Knockdown of rRNA Biogenesis Genes Protects Against Phenformin-Induced Reductive Stress

(A) Representative images of developmental assay performed on control *eft-3p::AtTIR-1(F79G)::mRuby;3xFLAG::AID*::GFP::FASN-1* animals fed RNAi and treated with a 0.1% DMSO or 5 µM 5-Ph-IAA diluted in either a water vehicle or 4.5 mM phenformin. Images were taken after 120 hours of treatment post-L1 of hatching.
